# *satuRn:* Scalable Analysis of differential Transcript Usage for bulk and single-cell RNA-sequencing applications

**DOI:** 10.1101/2021.01.14.426636

**Authors:** Jeroen Gilis, Kristoffer Vitting-Seerup, Koen Van den Berge, Lieven Clement

## Abstract

Alternative splicing produces multiple functional transcripts from a single gene. Dysregulation of splicing is known to be associated with disease and as a hallmark of cancer. Existing tools for differential transcript usage (DTU) analysis either lack in performance, cannot account for complex experimental designs or do not scale to massive scRNA-seq data. We introduce satuRn, a fast and flexible quasi-binomial generalized linear modelling framework that is on par with the best performing DTU methods from the bulk RNA-seq realm, while providing good false discovery rate control, addressing complex experimental designs and scaling to scRNA-seq applications.

## Introduction

Studying differential expression (DE) is one of the key tasks in the downstream analysis of RNA-seq data. Typically, DE analyses identify expression changes on the gene level. However, the widespread adoption of expression quantification through pseudo-alignment^1,2^, which enables fast and accurate quantification of expression at the transcript level, has effectively paved the way for transcript-level analyses. Here, we specifically address differential transcript usage (DTU) analysis, one type of transcript-level analysis that studies the change in relative usage of transcripts/isoforms within the same gene. DTU analysis holds great potential: previous research has shown that most multi-exon human genes are subject to alternative splicing and can thus produce a variety of functionally different isoforms from the same genomic locus^3–5^. The dysregulation of this splicing process has been reported extensively as a cause for disease^6–9^, including several neurological diseases such as frontotemporal dementia, Parkinsonism and spinal muscular atrophy, and is a well-known hallmark of cancer^10^.

In this context, full-length single-cell RNA-Seq (scRNA-seq) technologies such as Smart-Seq2^11^ and Smart-Seq3^12^ hold the promise to further increase the resolution of DTU analysis from bulk RNA-seq data towards the single-cell level, where differences in transcript usage are expected to occur naturally between cell types. However, only a few bespoke DTU methods have been developed for scRNA-seq data and they lack biological interpretation. Indeed, methods specifically developed for scRNA-seq data are either restricted to exon/event level^13,14^ analysis (e.g. pinpointing exons involved in splicing events), or they can only pinpoint DTU genes without unveiling the actual transcripts that are involved^15^. Interestingly, many DTU methods for bulk RNA-seq do provide inference at the transcript level and their performance has already been extensively profiled in benchmark studies^16–18^. Based on a subset of the simulated RNA-seq dataset from Love *et al*.^18^ (see Methods), we show the performance of six DTU tools; DEXSeq^19^, DoubleExpSeq^20^, DRIMSeq^21^, edgeR diffSplice^22^, limma diffSplice^23^ and NBSplice^24^ (Figure 1A). DEXSeq and DoubleExpSeq have a higher performance than the other methods. In addition, we observe that most methods, and DRIMSeq in particular, fail to control the false discovery rate (FDR) at its nominal level, which is in line with previous reports^16–18^.

**Figure 1:**
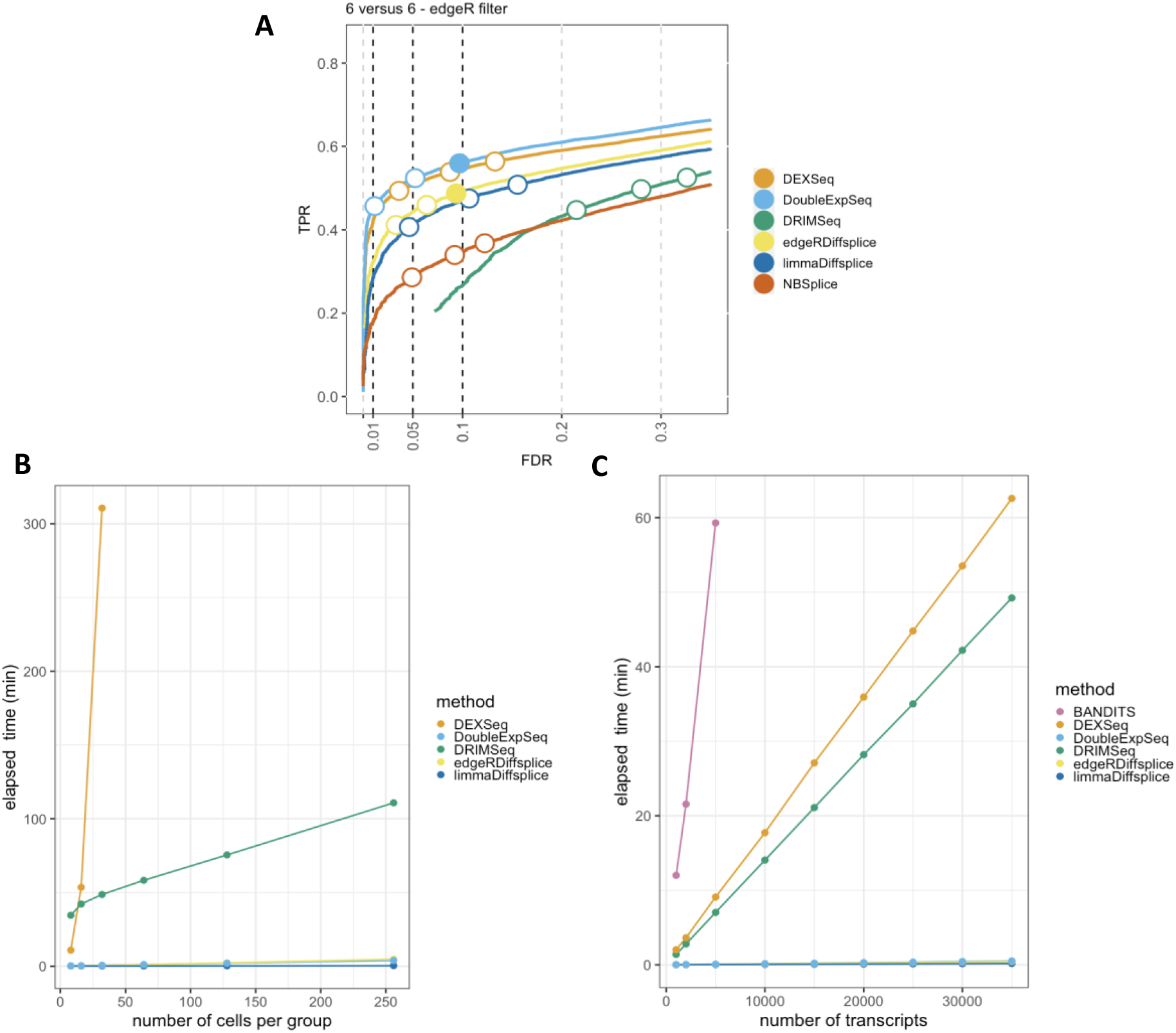
Performance and scalability evaluation of six DTU methods. **A: Performance evaluation on the simulated bulk RNA-Seq dataset from Love *et al***.^18^. Each curve displays the performance of each method by evaluating the sensitivity (TPR) with respect to the false discovery rate (FDR). The three circles on each curve represent working points when the FDR level is set at nominal levels of 1%, 5% and 10%, respectively. The circles are filled if the empirical FDR is equal or below the imposed FDR threshold. DEXSeq and DoubleExpSeq clearly have the highest performances. Note that most methods, and DRIMSeq in particular, fail to control the FDR at its nominal level. **B: Scalability with respect to the number of cells in a scRNA-Seq dataset**. While all other methods scale linearly with an increasing number of cells, DEXSeq scales quadratically. As such, DEXSeq cannot be used for the analysis of large bulk and scRNA-Seq datasets. For all sample sizes, the number of transcripts in the datasets were set at 30.000. Note that NBSplice needed to be omitted from this analysis as it fails to converge on datasets with a large proportion of zero counts (see below). **C: Scalability with respect to the number of transcripts in a scRNA-Seq dataset**. While all other methods scale linearly with an increasing number of cells, BANDITS scales quadratically. Moreover, BANDITS failed to run on our system for datasets with 7.500 transcripts or more. As such, it had to be omitted from panels A and B. A performance and scalability evaluation of BANDITS on datasets with an (artificial) lower number of transcripts is provided in supplementary Figures S1 and S3.

In order to assess DTU in single-cell applications, however, these bulk RNA-seq DTU tools should scale to the large data volumes generated by full-length scRNA-seq platforms, which can profile the transcriptome of several thousands of cells^25–27^ in a single experiment. In Figure 1B, we evaluate the required computational time in function of the number of sequenced libraries for a two-group DTU analysis for 30,000 transcripts on a subset of the scRNA-seq dataset from Chen *et al*.^28^. In spite of its good performance, the popular tool DEXSeq already required more than five hours to analyze two groups of 32 cells and clearly does not scale to large bulk nor scRNA-seq datasets.

In addition, DTU methods should allow for the analysis of datasets with large numbers of (unique) transcripts. The number of transcripts that are typically assessed depends on the coverage of the RNA-seq experiment and the adopted filtering criteria in the data analysis workflow. As the coverage of RNA-seq experiments has increased rapidly over the past few years and can be expected to continue expanding, scalability towards large numbers of transcripts will be essential to enable a transcriptome-wide view on the isoform usage changes. In Figure 1C, we perform a DTU analysis across a range of transcripts in a two-group comparison with 16 cells each, using the dataset from Chen *et al*. Here, we observed that the DTU tool BANDITS^29^ scales particularly poorly to large numbers of transcripts. More specifically, BANDITS did not complete the DTU analysis on the dataset with 7.500 transcripts within 137 hours on our system (see Methods); therefore, larger analyses were omitted. As such, BANDITS had to be omitted from the analyses shown in Figures 1A and 1B. For a performance and scalability evaluation of BANDITS on datasets with an (artificial) lower number of transcripts, we refer to Figures S1 and S3.

Besides scalability, several other issues arise when porting bulk RNA-seq DTU tools towards scRNA-seq applications. Indeed, modeling scRNA-seq data often requires multifactorial designs, for instance when comparing expression levels across multiple cell types between multiple treatment groups. Accounting for multiple covariates, however, is not implemented in BANDITS, NBSplice and DoubleExpSeq, jeopardizing their utility for (sc-)RNA-seq DTU analysis. Another issue arises with the large numbers of zero counts in scRNA-seq data, which seems to be particularly problematic for NBSplice that fails to converge if the gene-level count of any of the samples or cells is zero. As such, NBSplice could not be evaluated in Figures 1B and 1C.

Altogether, many of the existing DTU analysis tools are not well suited to analyze large bulk RNA-seq and full-length scRNA-seq datasets, leaving the great potential of these data largely unexploited. In light of these shortcomings we developed *satuRn*, which is an acronym for Scalable Analysis of differential Transcript Usage for RNa-seq data, a novel method for DTU analysis that (i) is highly performant, (ii) provides a good control of the false discovery rate (FDR) (iii) scales seamlessly to the large data volumes of contemporary (sc-)RNA-seq datasets, (iv) allows for modelling complex experimental designs, (v) can deal with realistic proportions of zero counts and (vi) provides direct inference on the biologically relevant transcript level. In brief, satuRn adopts a quasi-binomial (QB) generalized linear model (GLM) framework. satuRn provides direct inference on DTU by modelling the relative usage of a transcript, in comparison to other transcripts from the same gene, between groups of interest. To stabilize the estimation of the overdispersion parameter of the QB model, we borrow strength across transcripts by building upon the empirical Bayes methodology as introduced by Smyth *et al*.^23^. In order to control the number of false positive findings, an empirical null distribution is used to obtain the p-values, which are corrected for multiple testing with the FDR method of Benjamini and Hochberg^30^. Our method is implemented in an R package available at https://github.com/statOmics/satuRn and will be submitted to the Bioconductor project.

## Results

We first evaluate the performance of our novel DTU method, satuRn, on publicly available simulated and real bulk RNA-seq data, as well as on real scRNA-seq data. In general, we found that the performance of satuRn was at least on par with the performances of the best tools from the literature. In addition, our method controls the FDR closer to the nominal level, on average. Second, we show that satuRn scales towards the large data volumes generated by contemporary bulk and single-cell RNA-seq experiments, allowing for a transcriptome-wide analysis of datasets consisting of several thousands of cells, in only a few minutes. Finally, we analyze a large full-length scRNA-seq case study dataset, where we obtain highly relevant biological results on isoform-level changes between cell types that would have remained obscured in a canonical differential gene expression (DGE) analysis.

### Performance on simulated bulk RNA-seq datasets

To evaluate the performance of satuRn, we adopt three simulated bulk RNA-seq datasets from previous publications. Dataset 1 was obtained from Love *et al*.^18^ and contains two groups of twelve samples each, which we subsample without replacement to evaluate 3vs3, 6vs6 and 10vs10 two-group comparisons. Datasets 2 and 3 are the *Drosophila melanogaster* and *Homo sapiens* simulation studies from Van den Berge *et al*.^31^ and Soneson *et al*.^17^, which both contain two groups of five samples each. In brief, all datasets were constructed by generating sequencing reads based on parameters that are estimated from real bulk RNA-seq data. DTU between groups of samples was artificially introduced in the data, prior to the quantification of expression using either Salmon^2^ (dataset 1) or kallisto^1^ (dataset 2 and 3). Notably, there are some methodological differences between the simulation framework of dataset 1 and that of datasets 2 and 3 with respect to the read generation and the simulation of DTU signal (see Methods). In terms of transcript filtering, we adopt two different strategies as implemented by edgeR^32^ and DRIMSeq^21^, which correspond to a lenient and more stringent filtering, respectively (see Methods).

The result of the performance evaluation of satuRn with respect to other DTU methods on the three simulated bulk datasets is displayed in Figure 2. Figure 2A shows the average performance over three *6 versus 6* subsamples for dataset 1, after filtering with edgeR. Figures 2B and 2C display the performance on datasets 2 and 3 after edgeR filtering, respectively. In all three datasets, satuRn outperforms NBSplice, edgeR diffsplice and limma diffsplice. Intriguingly, the performance of DRIMSeq varies strongly between the three datasets. This discrepancy may be explained by the different strategies for generating reads and introducing DTU between dataset 1, and, datasets 2 and 3 (see methods). We furthermore find the performance of satuRn is on par with the best performing tools from the literature, DEXSeq and DoubleExpSeq. In addition, both satuRn and DoubleExpSeq provide a stringent control of the FDR, while DEXSeq and DRIMSeq are often too liberal, as reported previously^18^.

**Figure 2:**
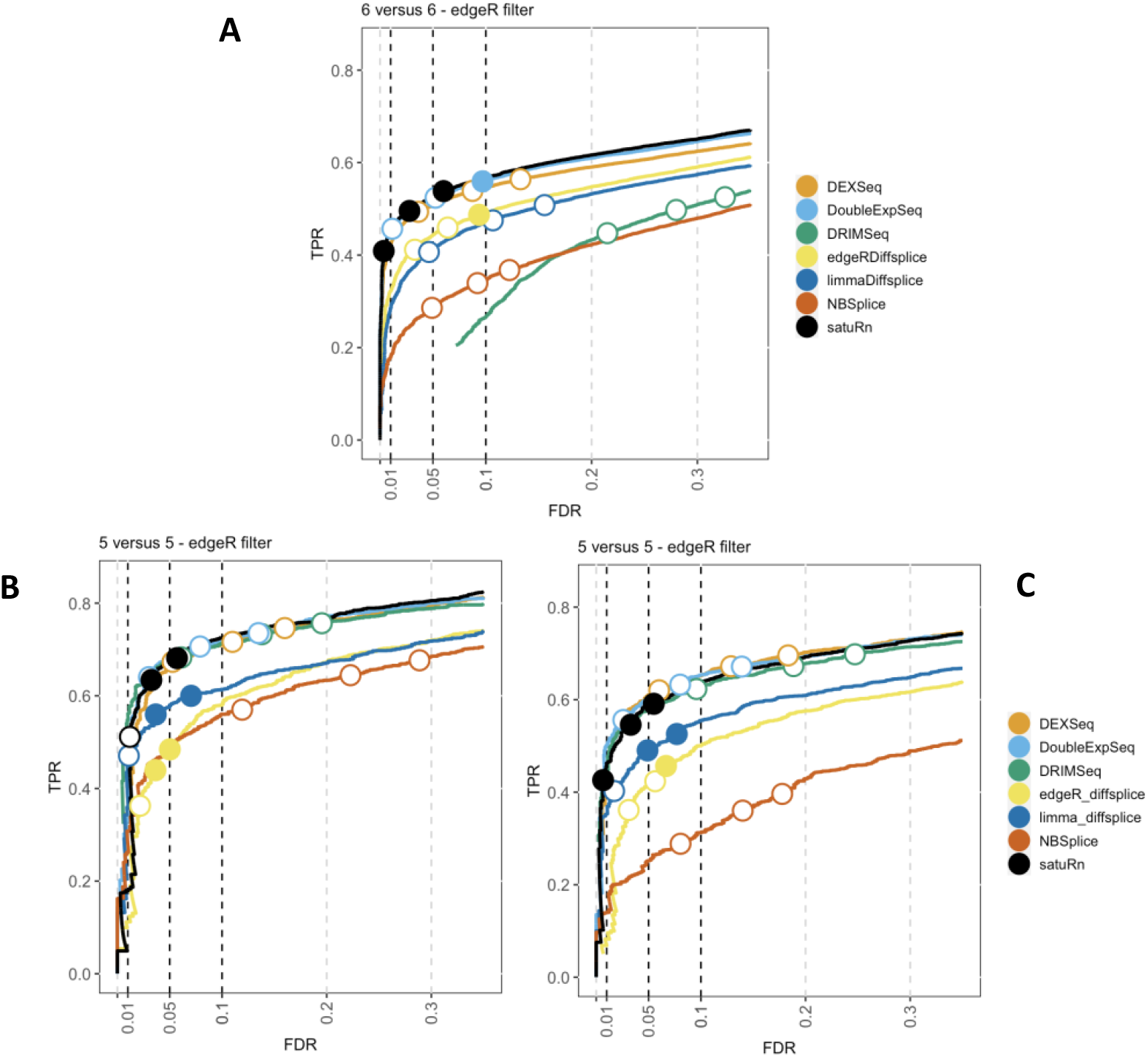
Performance evaluation of *satuRn* on three simulated bulk RNA-Seq datasets. Each curve visualizes the performance of each method by displaying the sensitivity of the method (TPR) with respect to the false discovery rate (FDR). The three circles on each curve represent working points when the FDR level is set at nominal levels of 1%, 5% and 10%, respectively. The circles are filled if the empirical FDR is equal or below the imposed FDR threshold. The performance of *satuRn* is on par with the best tools from the literature, DEXSeq and DoubleExpSeq, for all datasets. In addition, our method consistently controls the FDR close to its imposed nominal FDR threshold.

We also evaluated the effects of sample size and different filtering criteria on the performance of the different DTU methods (see Figures S2, S3, S4 and S5). Neither sample size nor filtering criterion had a profound impact on the ranking of the performances of the different DTU methods; satuRn, DEXSeq and DoubleExpSeq remain the best performing methods overall. In addition, we studied the impact of using either raw count estimates or normalized abundance estimates (scaledTPM, see Methods) as input data for the DTU algorithms. We observed a slightly higher performance in all datasets when providing raw abundance estimates, except for Dataset 1 from Love *et al*.^18^. All performance evaluations in the body of this publication therefore were generated with raw count estimates as input data, except for Figure 2, panel A. For a full overview on the effects of sample size, filtering criteria and data input type, we refer to supplementary figures S2-S9.

### Performance on a real bulk RNA-seq dataset

While simulation studies are common for evaluating the performance of DE analysis methods, there is currently no consensus on the simulation strategy that best mimics real (sc)RNA-seq data. In addition, simulation frameworks typically generate data according to parametric assumptions on the data-generating mechanism, thus potentially favoring DE methods that adopt similar distributional assumptions in their statistical model^33^. An alternative procedure is to non-parametrically modify a real dataset. Here, we obtained different subsamples from the large bulk RNA-seq dataset available from the Genotype-Tissue Expression (GTEx) consortium^34^, generating 9 datasets in total, i.e. 3 repeats for each of 3 sample sizes; *5 versus 5, 20 versus 20* and *50 versus 50* samples. We then artificially introduced DTU in these data by swapping transcript usages between groups of samples (see Methods for details). Again, we adopt two different filtering strategies as implemented by edgeR^32^ and DRIMSeq^21^ (see Methods).

The results of the performance evaluation of satuRn on the real bulk datasets upon edgeR filtering is displayed in Figure 3. In agreement with the results obtained from the simulated bulk RNA-seq study, we observe that the performance of satuRn is on par with DEXSeq and DoubleExpSeq. Again, satuRn provides a conservative FDR control. While the FDR control of DoubleExpSeq is good overall, it appears to become too liberal with increasing sample size. In this evaluation, DRIMSeq performs poorly, in contrast to simulated bulk RNA-seq datasets 2 and 3, but in line with the performance evaluation on the simulated bulk RNA-seq dataset 1. Note that DEXSeq, DRIMSeq and NBSplice were omitted from the analysis of the largest dataset (*50 versus 50* samples), as these methods do not scale to such large datasets (Figure 1). Adopting the DRIMSeq-based filtering did not have a qualitative impact on the performance (Figure S6).

**Figure 3:**
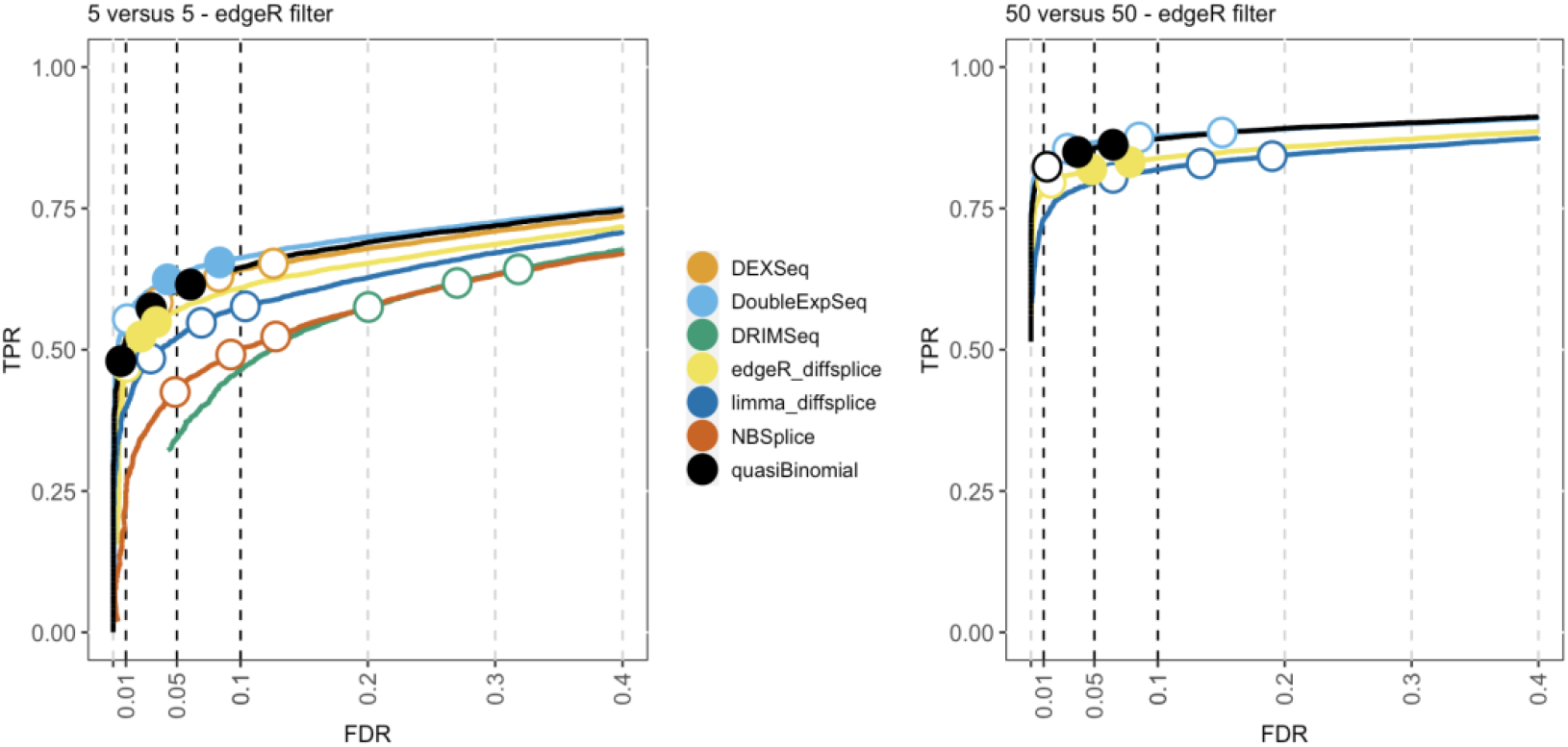
Performance evaluation of satuRn on a real bulk RNA-Seq dataset. Each curve visualizes the performance of each method by displaying the sensitivity of the method (TPR) with respect to the false discovery rate (FDR). The three circles on each curve represent working points when the FDR level is set at nominal levels of 1%, 5% and 10%, respectively. The circles are filled if the empirical FDR is equal or below the imposed FDR threshold. The performance of satuRn is on par with the best tools from the literature, DEXSeq and DoubleExpSeq. In addition, satuRn consistently controls the FDR close to its imposed nominal FDR threshold, while DoubleExpSeq becomes more liberal with increasing sample sizes. Note that DEXSeq, DRIMSeq and NBSplice were omitted from the larger comparison, as these methods do not scale to large datasets (Figure1).

### Performance on real single-cell data

Finally, we evaluate the performance of satuRn on single-cell RNA-seq data. As with the real bulk analysis, the single-cell datasets were generated by subsetting from three different real scRNA-seq datasets^25,28,35^ (see Methods). Again, we subsampled three repeats of different sample sizes, artificially introduced DTU with the swapping strategy and applied either the edgeR- or DRIMSeq-based filtering criterium (see Methods for details).

By subsampling the Chen et al.^28^ dataset, we generated three repeats of two sample sizes, i.e. *20 versus 20* and *50 versus 50 cells*. The results of the performance evaluation of satuRn on this dataset upon edgeR filtering is displayed in Figure 4. The performance of satuRn is slightly better than that of the best tool from the literature, DoubleExpSeq. As compared to the evaluations on bulk data, we observe a performance drop for DEXSeq relative to satuRn and DoubleExpSeq. This, in combination with its poor scalability (Figure 1), greatly compromises the use of DEXSeq for the analysis of scRNA-seq data. satuRn again provides a stringent control of the FDR, while the inference of DoubleExpSeq is too liberal, again becoming more problematic for larger sample sizes. Adopting the DRIMSeq filter did not have a qualitative impact on the performances (Figure S7). The results of the performance evaluations on the other two scRNA-seq datasets^25,35^ are in strong agreement with the results displayed here, with satuRn performing at least on par with DoubleExpSeq and satuRn additionally controlling the FDR around the nominal level (Figures S8 and S9).

**Figure 4:**
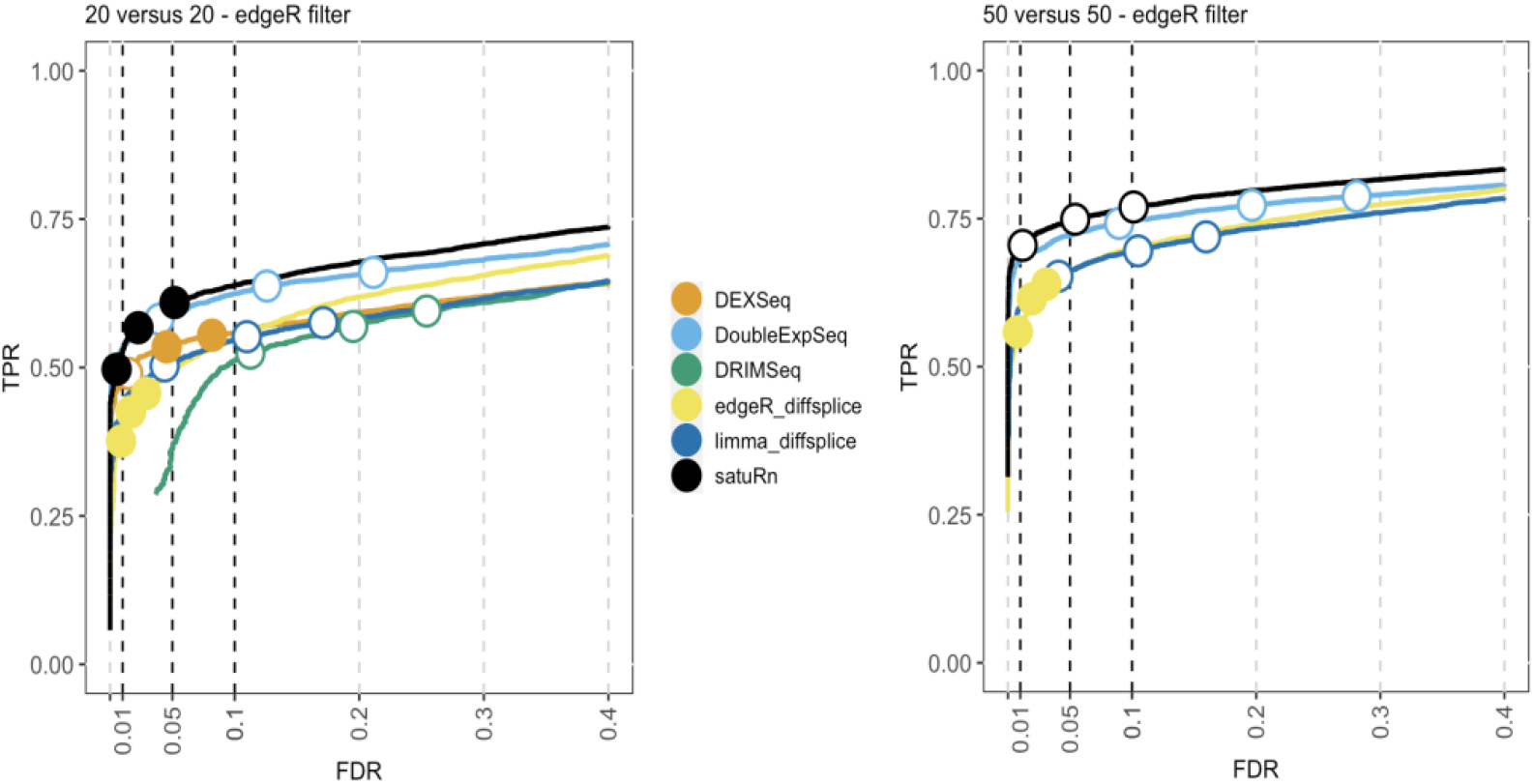
Performance evaluation of satuRn on a real scRNA-Seq dataset. Each curve visualizes the performance of each method by displaying the sensitivity of the method (TPR) with respect to the false discovery rate (FDR). The three circles on each curve represent working points when the FDR level is set at nominal levels of 1%, 5% and 10%, respectively. The circles are filled if the empirical FDR is equal or below the imposed FDR threshold. The performance of satuRn is on par with the best tools from the literature, DEXSeq and DoubleExpSeq. In addition, our method provides a stringent control of the FDR, while DoubleExpSeq becomes more liberal with increasing sample sizes (see also Figure S6). Note that DEXSeq and DRIMSeq were omitted from the two largest comparisons, as these methods do not scale to large datasets (Figure1). NBSplice was omitted from all comparisons, as it does not converge on datasets with many zeros, such as scRNA-Seq datasets.

Notably we found that the theoretical null distribution of the test statistics from satuRn failed to provide good FDR control in single-cell analyses (Figure S10). To obtain proper p-values with satuRn in single-cell applications, we therefore estimate the null distribution of the test statistic empirically (see Methods, satuRn paragraph). Note, that the use of the empirical null distribution in our bulk RNA-seq benchmarks does not affect the results because no deviations of the theoretical null distribution occur. However, the empirical null resulted in much improved FDR control in scRNA-seq datasets (Figure S10). We therefore adopt the empirical null estimation as the default setting in satuRn. As such, all satuRn results in this publication are relying on the empirical null strategy. As a final remark, we likewise attempted to improve the FDR control of DoubleExpSeq. However, in all analyses with DoubleExpSeq we observed a large spike of p-values equal to 1, which poses a problem when estimating the empirical null distribution (Figure S11). Therefore, this strategy could not be used to improve the FDR control of DoubleExpSeq.

### Scalability benchmark

We performed a computational benchmark of satuRn to investigate its scalability with respect to the number of samples/cells and the number of transcripts in an RNA-seq dataset. All scalability benchmarks were run on a single core of a Linux machine with an Intel(R) Xeon(R) CPU E5-2420 v2 (2.20GHz, Speed: 2200 MHz) processor and 30GB RAM. The results are displayed in Figure 5.

**Figure 5:**
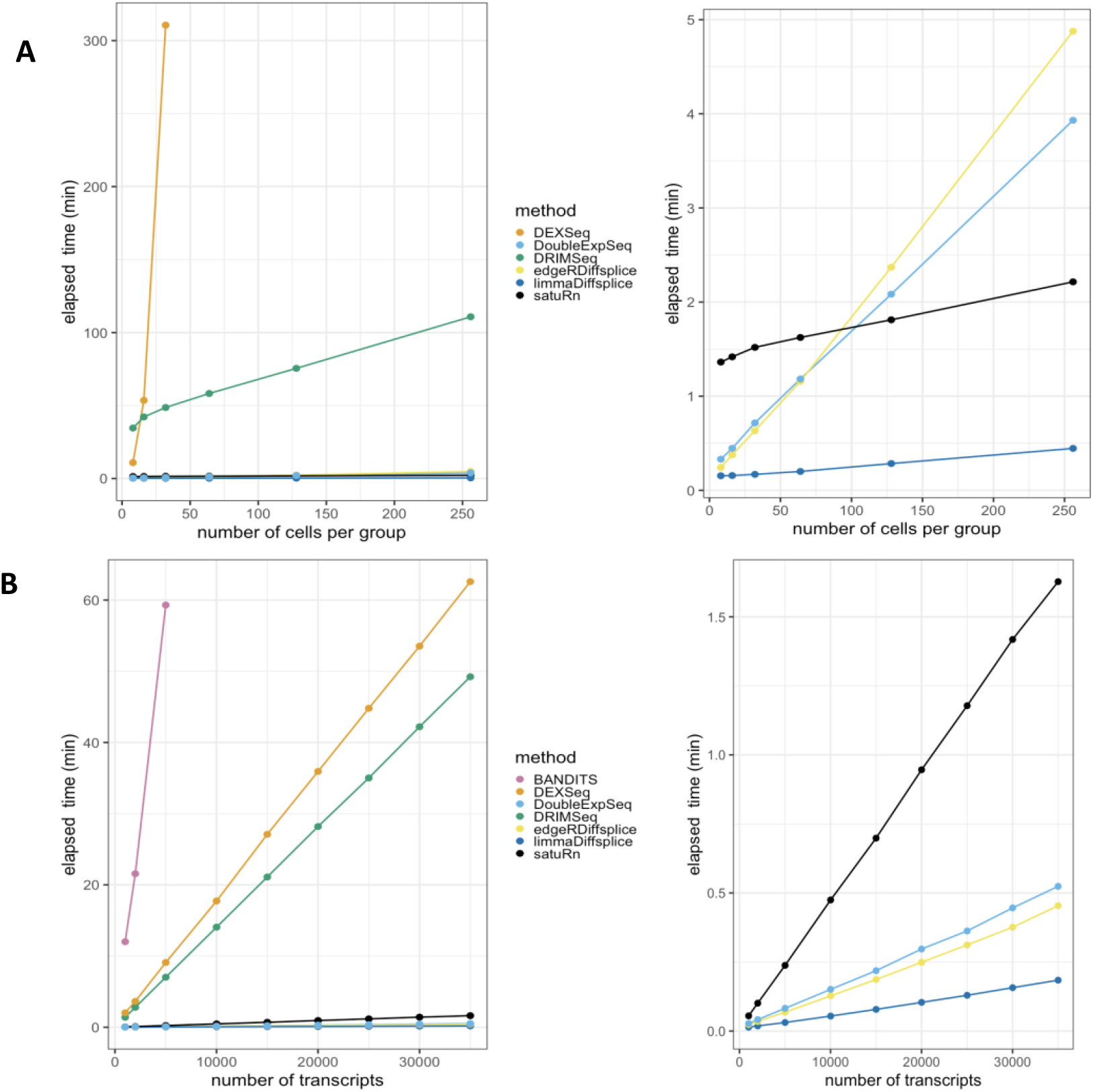
Scalability evaluation of satuRn on scRNA-Seq data. **A: Runtime with respect to the number of cells in a scRNA-Seq dataset. Left panel:** it is clear that DRIMSeq and especially DEXSeq scale very poorly with the number of cells in the dataset. **Right panel:** Detailed plot of the remaining methods. satuRn scales linearly with increasing numbers of cells, with a slope that is comparable to that of limma diffsplice. As such, satuRn is able to perform a DTU analysis on a dataset with two groups of 256 cells each and 30.000 transcripts in less than three minutes. For all sample sizes, the number of transcripts in the datasets were set at 30.000. Note that NBSplice was not included in this analysis as it fails to converge on datasets with a large proportion of zero counts. **B: Runtime with respect to the number of transcripts in a scRNA-Seq dataset. Left panel:** DEXSeq, DRIMSeq and especially BANDITS scale poorly to the number of transcripts in the dataset. **Right panel:** Detailed plot of the remaining methods. satuRn scales linearly with increasing numbers of transcripts, but with a steeper slope than edgeR diffsplice, DoubleExpSeq and limma diffsplice. The number of cells in the dataset was set fixed to two groups of 16 cells. All scalability benchmarks were run on a single core.

Figure 5A displays the scalability with respect to the number of cells in the dataset, while keeping the number of transcripts in the dataset fixed at 30.000. From the left panel, it is clear that DRIMSeq and especially DEXSeq scale very poorly with the number of cells in the dataset, which was already shown in Figure 1B. In the right panel, we focus on the four remaining methods. satuRn scales linearly with increasing numbers of cells, with a slope comparable to limma diffsplice. As such, satuRn is able to perform a DTU analysis on a dataset with two groups of 256 cells each and 30.000 transcripts in less than three minutes. Note that BANDITS^29^ was not included in this benchmark, as it does not scale to datasets with this many transcripts. For a performance and scalability evaluation of BANDITS on datasets with a lower number of transcripts, we refer to Figure S1. NBSplice was also omitted as it fails to converge on datasets with a large proportion of zero counts.

Figure 5B shows the scalability with respect to the number of transcripts in the dataset, while keeping the number of cells in the dataset fixed to two groups of 16 cells. As shown in Figure 1C, BANDITS, DEXSeq and DRIMSeq scale poorly to datasets with many transcripts. From the right panel, satuRn scales linearly with increasing numbers of transcripts, albeit with a steeper slope than edgeR diffsplice, DoubleExpSeq and limma diffsplice. Note, that the scalability of DTU analyses can be improved through parallelization, if this is allowed by the underlying algorithm. Parallel execution is implemented in satuRn and in all methods from the literature that were discussed in this manuscript, except for DoubleExpSeq and NBSplice.

Altogether, we find that while several methods for DTU analysis exist, none are optimally suited for analyzing single-cell datasets. DRIMSeq, NBSplice, edgeR diffsplice and limma diffsplice have a much lower overall performance in our benchmarks. DEXSeq does not scale to large datasets. Finally, DoubleExpSeq does not support experimental designs that require an analysis with multiple additive effects, e.g. randomized complete block designs and designs where batch-effect correction is required, which are essential for many practical scRNA-Seq analysis settings^36^. In addition, it fails to control the FDR at the desired level, especially with increasing sample sizes.

### Case study

We use satuRn to perform a DTU analysis on a subset of the single-cell (SMART-seq2^11^) RNA-seq dataset from Tasic *et al*.^35^. In addition, we analyze the same dataset with DoubleExpSeq and limma diffsplice, which are the only other DTU methods that scale to large scRNA-seq datasets and have a reasonable performance in our benchmarks. In the original publication, the authors studied differential gene expression between cell types originating from two areas at distant poles of the mouse neocortex; the primary visual cortical area (VISp), which processes sensory information with millisecond timescale dynamics^37–39^ and the anterior lateral motor cortex (ALM), which displays slower dynamics related to short-term memory, deliberation, decision-making and planning^40,41^. Based on marker genes, Tasic *et al*.^35^ assigned all of the 23.822 cells from the scRNA-seq dataset to one of three cell classes; glutamatergic (excitatory) neurons, GABAergic (inhibitory) neurons or non-neuronal cells. The authors then further classified the neuronal cells into several subclasses based on their dominant layer of dissection and projection patterns (through a retrograde labelling experiment). Finally, these subclasses are further classified into cell types based on the expression of specific marker genes.

### DGE analysis with edgeR

In their original DGE analysis, Tasic *et al*.^35^ obtained the largest number of differentially expressed genes between the cell types originating from the ALM and VISp regions of the glutamatergic L5 IT subclass (2.739 cells in total), where L5 refers to layer-of-dissection 5 and IT refers to the intratelencephalic projection type. Here, we first perform a DGE analysis with an edgeR-based workflow (see Methods) on the same comparisons between L5 IT cell types that were assessed by Tasic *et al*. Table 1 shows the number of differentially expressed genes between the groups of interest in column 4.

**Table 1:**
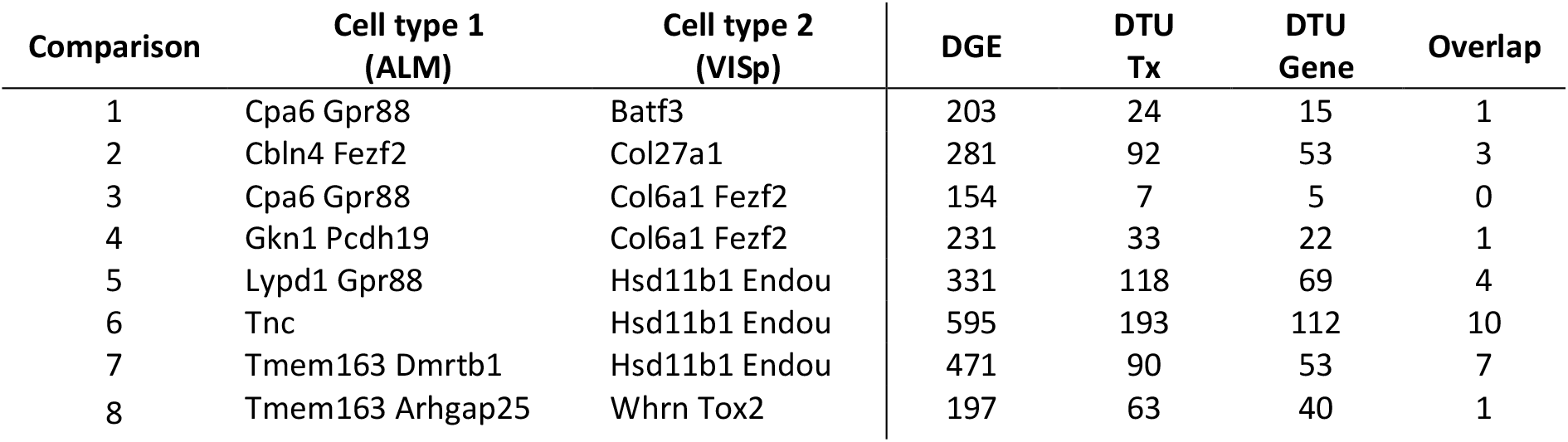
Number of differentially expressed genes and differentially used transcripts in eight comparisons between cell types. The first three columns indicate the comparisons between ALM (column 2) and VISp (column 3) cell types, respectively. Column 4 indicates the number of differentially expressed genes as identified with an edgeR analysis. Column 5 displays the number of transcripts that satuRn flagged as differentially used. Column 6 shows the number of unique genes in which satuRn finds evidence of differential usage of at least one transcript. Column 7 displays the absolute number of genes that overlap between columns 4 and 6.

### DTU analysis with satuRn

Next, we perform a DTU analysis for the same cell types using satuRn. In column 5 of Table 1, we display the number of differentially used transcripts for each comparison. We also show the number of unique genes in which we find evidence for changes in usage of at least one transcript (column 6). While the number of differentially used transcripts is lower than the number of differentially expressed genes in each of the contrasts, we did identify differentially used transcripts in all contrasts of interest. Most interestingly, we observe that the overlap between the differentially expressed genes and the genes in which we found evidence for DTU is very limited (Table 1, column 7). This shows that the information obtained from our DTU analyses are orthogonal to the results from the canonical DGE analyses, which has been reported previously for simulated bulk data^18^.

#### Gene set enrichment analysis

We perform a gene set enrichment analysis (GSEA, see Methods) on the three comparisons with most DE genes and most genes with evidence for DTU (comparisons 5, 6 and 7). Similar gene ontology categories are returned for the set of DGE genes and the set of DTU genes, with many of the enriched processes being biologically relevant in the context of this case study. Enriched gene sets include the gene ontology classes, synapse, neuron projection, synaptic signaling and cell projection organization. This shows that the complementary information brought by the DTU analysis is indeed biologically relevant. For an extensive overview of the GSEA of the set of DGE genes and genes with evidence of DTU in comparisons 5, 6 and 7, we refer to supplementary table 1.

#### satuRn identifies biologically relevant DTU transcripts

To display the utility of satuRn for identifying and visualizing DTU transcripts in scRNA-seq datasets, we focus on comparison #6 of the DTU analysis and discuss the gene P2X Purinoceptor 4 or P2rx4 (Ensembl ID ENSMUSG00000029470), a gene which is part of a family of purinergic receptors that have been implicated in functions such as learning, memory and sleep. In the DGE analysis, no evidence for differential expression of P2rx4 was found at the gene level (FDR-adjusted p-value = 1). By contrast, in our DTU analysis the transcripts of P2xr4 displayed the highest statistical evidence for differential usage within the set of transcripts that could be assigned to the ontology class ‘neuron projection’^42^. The mean usage of transcript ENSMUST00000081554 is estimated to be 28.9% in Tnc cells and 75.9% in Hsd11b1 Endou cells (FDR-adjusted p-value = 1.22E-13). For transcript ENSMUST00000195963, the transcript usage changes from 58.2% in Tnc cells and 16.6% in Hsd11b1 Endou cells (FDR-adjusted p-value = 1.79E-10). For the third transcript of P2rx4 that was assessed in our DTU analysis, ENSMUST00000132062, we found no statistical evidence for DTU (FDR-adjusted p-value = 0.534). In Figure 6, we show the output for the visualization of the transcript usages for P2rx4 as obtained from satuRn. Interestingly, the majority transcript in the Tnc cell type, ENSMUST00000195963, is not protein coding^43^. By contrast, the majority transcript in the Hsd11b1 Endou cell type, ENSMUST00000081554, is coding for the P2X purinoceptor protein (UniProt ID Q9Z256). As such, the changes in transcript usage between both cell types represent actual biological differences in the functionality of the gene products, which may be relevant to the process of neuron projection. This functional difference would have remained obscured when only performing a canonical DGE analysis.

**Figure 6:**
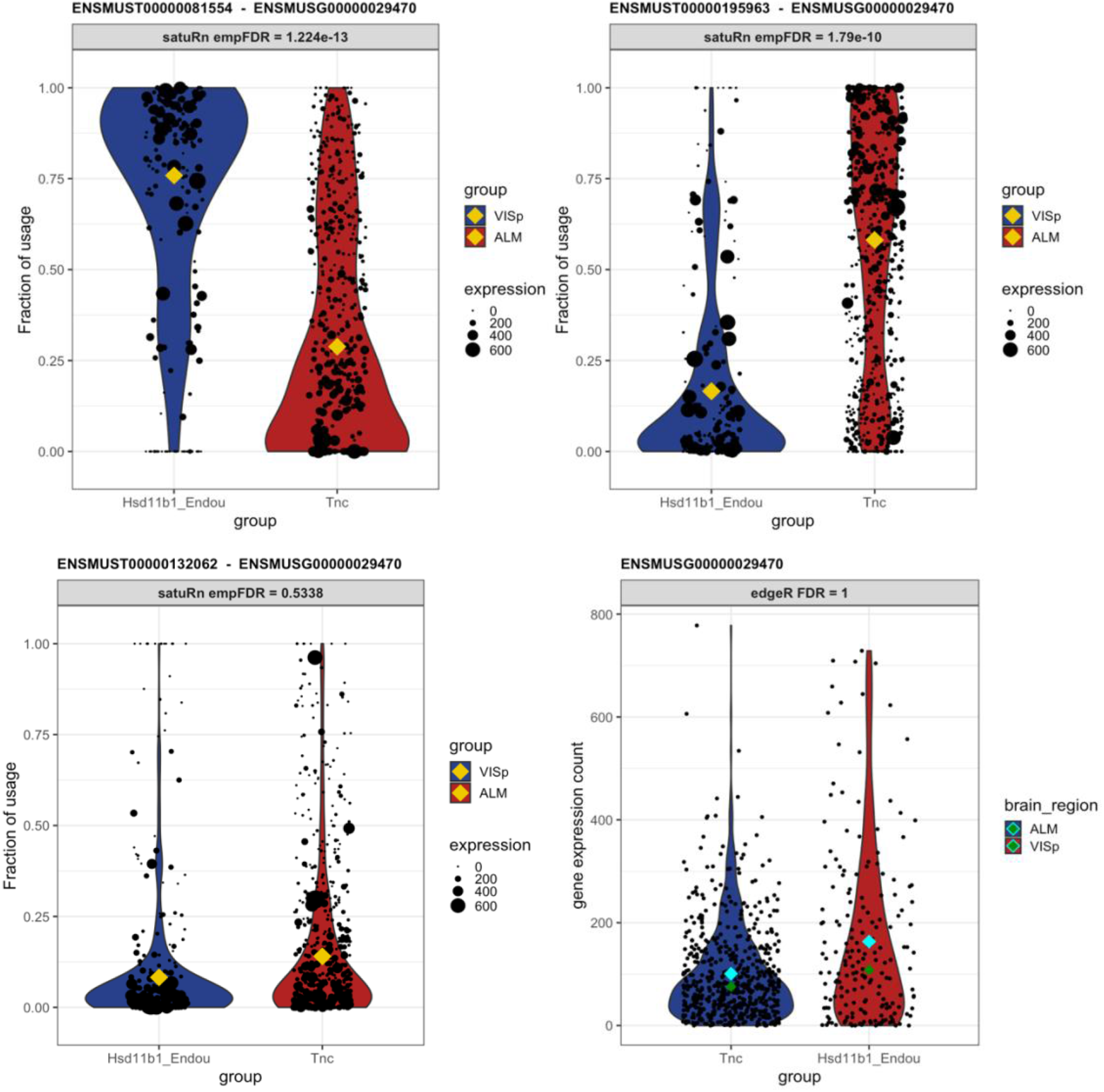
Differential transcript usage in the P2rx4 gene. Each panel shows transcript usage or gene expression across cells of the Tnc and Hsd11b1 cell types. For the transcript-level figures, the size of each datapoint is weighted according to the total expression of the gene in that cell, i.e. the gene counts per cell. The yellow diamonds indicate the estimated mean usage of a transcript for each cell type, as estimated by satuRn. The cyan and dark green diamonds indicate mean and median gene expression levels per cell type, respectively. The two top panels display the transcript usage across cells of the Tnc and Hsd11b1 Endou cell types of transcripts ENSMUST00000081554 and ENSMUST00000195963, respectively. The proportion of usage of the former transcript is clearly higher in Hsd11b1 Endou cells, while the latter transcripts is most abundant in Tnc cells. For the third transcript, ENSMUST00000132062 (bottom left panel) there is no evidence for differential usage between both cell types. In addition, there is no evidence for differential expression of P2rx4 on the gene level (bottom right panel). DTU and DGE significance levels are indicated in the figure headers.

### Comparison to limma diffsplice

We also analyzed the case study dataset with limma diffsplice^23^. When running limma diffsplice with default settings, a large number of DTU transcripts was returned (Figure S12) and we observe that the p-values were shifted towards smaller values (Figures S13 and S14). Therefore, we adopted the same empirical null strategy as implemented in satuRn to post-process the results. While this dramatically decreased the number of significant DTU transcripts, limma diffsplice still identified more transcripts (i.e. true or false positives) than our method. However, when we inspected the transcripts that were highly ranked in the top DTU list of limma diffsplice but lowly ranked in our top list, we found that most of these transcripts either originate from genes that are lowly expressed, or they are transcripts with a large fraction of zero counts (i.e. zero expression in a large percentage of cells). Limma diffsplice thus claims differential usage more often for transcripts that only contain little information for assessing DTU. This is depicted in Figure 7.

**Figure 7:**
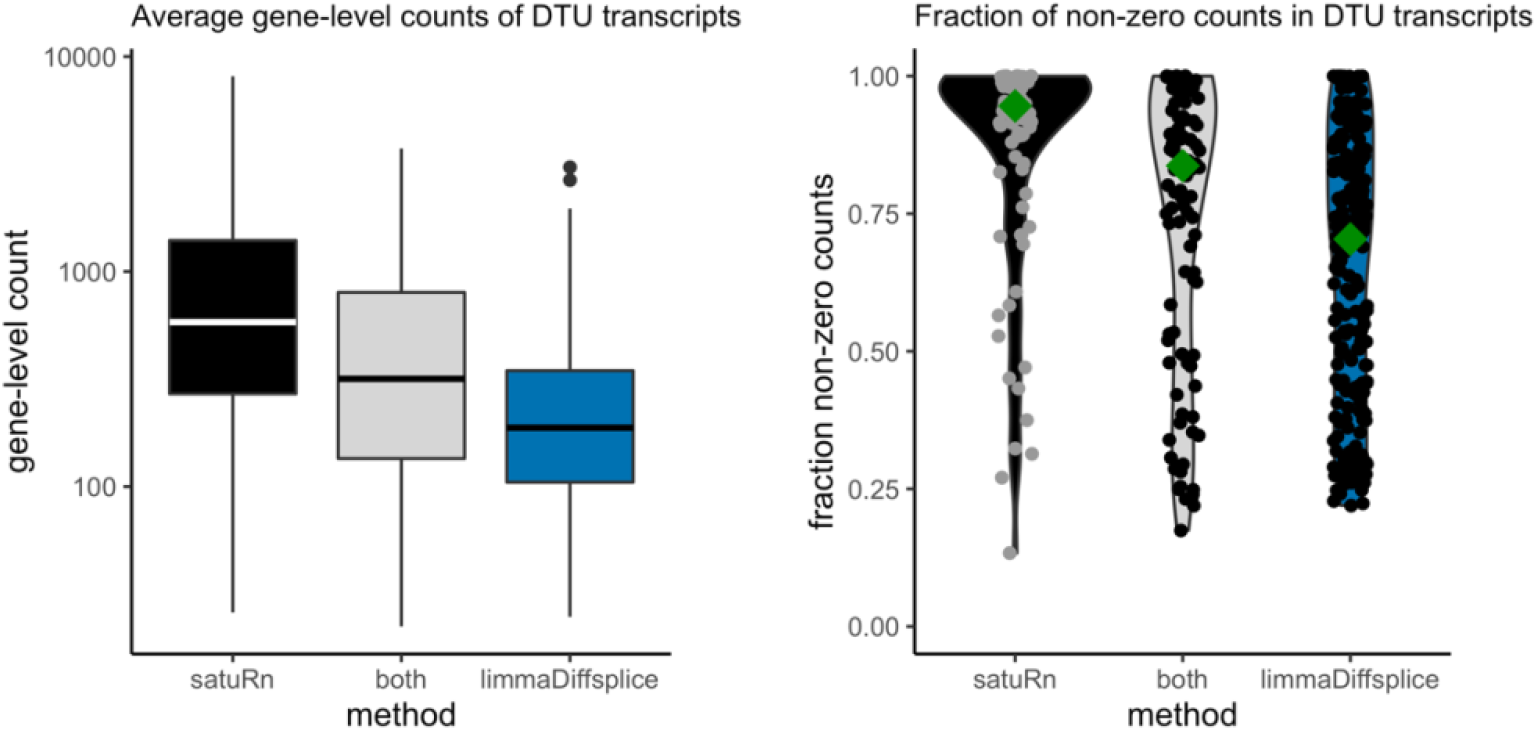
Global comparison between DTU transcripts uniquely identified by satuRn, uniquely identified by limma diffsplice or by both methods. **Left panel:** Boxplots on the average gene-level count for the DTU genes identified by the respective methods. Transcripts uniquely identified by satuRn originate from genes that have a much higher gene-level count (averaged over cells) as compared to transcripts uniquely identified by limma diffsplice. Note that the y-axis is displayed on a log10 scale. **Right panel:** Violin plots indicating the fraction of cells in which the transcripts are expressed. Transcripts uniquely identified by satuRn are expressed, on average, in a much larger fraction of the cells. Conversely, transcripts identified as DTU uniquely by limma diffsplice often have no expression in a large fraction of the cells. The dark green diamond indicates the median fraction of cells in which the DTU transcripts are expressed.

This behavior can be expected. Limma diffsplice tests for DTU by comparing the log-fold change in expression of transcript *t* with the average log-fold change in the expression of all transcripts belonging to the same gene as transcript *t*. As such, limma diffsplice does not incorporate any information on the absolute gene expression levels. In contrast, our quasi-binomial GLM framework models the log-odds of drawing a particular transcript *t* from the pool of transcripts in the corresponding gene. As a consequence, transcripts belonging to lowly expressed genes are correctly considered less informative in satuRn and are thus less likely to be picked up. For example, in Figure 8A, we show that while our method estimates a mean usage of 7% in Tnc cells and 26% in Hsd11b1 Endou cells (indicated by the gold diamond), the transcript is not identified as differentially used, given the low abundance of the corresponding gene and the highly variable single-cell level observations.

**Figure 8:**
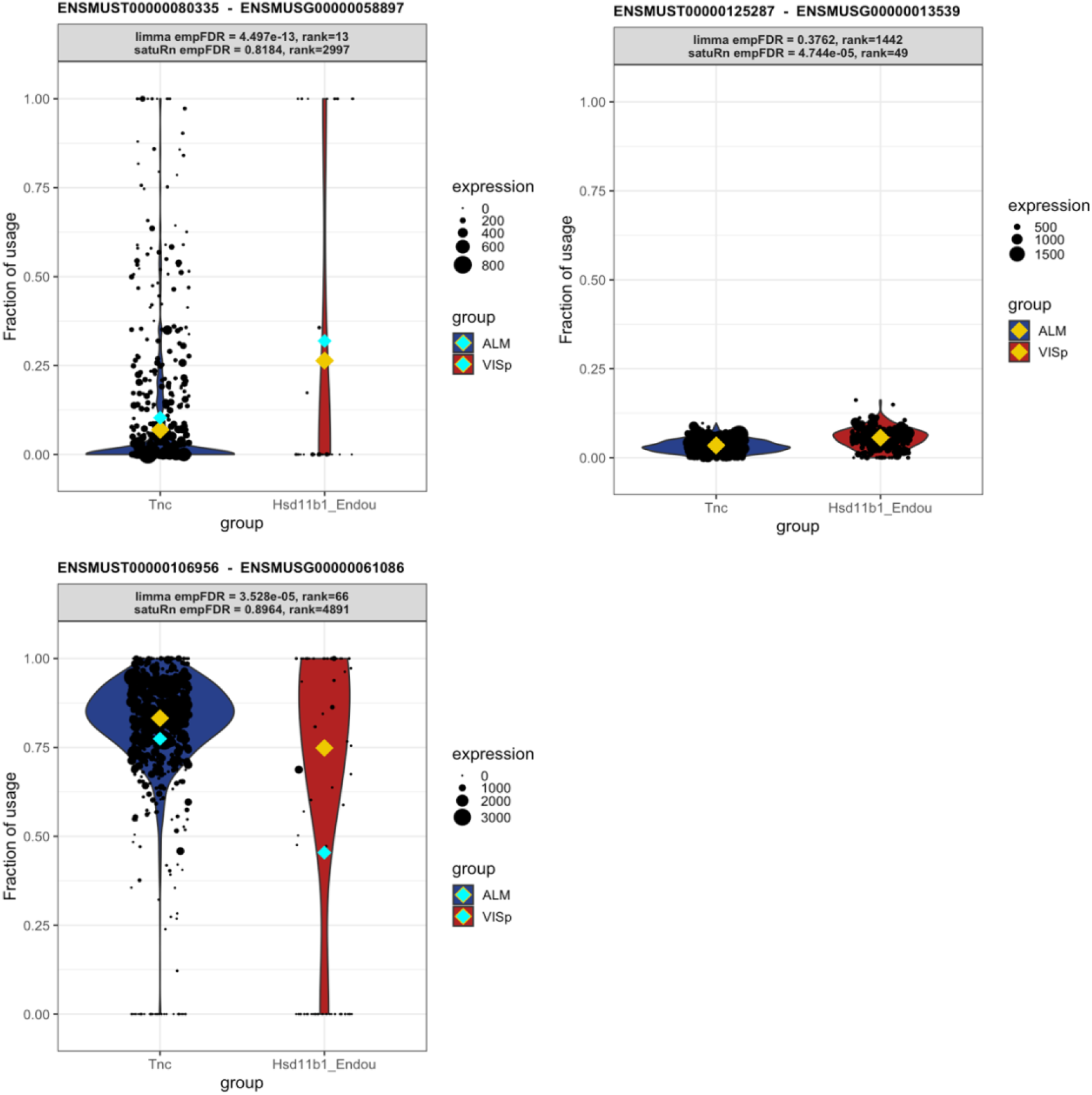
Three examples displaying DTU transcripts that are uniquely identified by satuRn or limma diffsplice. Each panel shows transcript usage across cells of the Tnc and Hsd11b1 cell types. The size of each datapoint is weighted according to the total expression of the corresponding gene in that cell, i.e. the total gene count per cell. The yellow diamonds indicate the estimated mean usage of a transcript for each cell type, as estimated by satuRn. The cyan diamonds indicate the mean transcript expression levels per cell type. The header of each panel indicates the FDR-adjusted p-value and the rank of the DTU finding in the top lists by limma diffsplice and satuRn analyses. **Panel A:** Transcript uniquely identified as differentially used by limma diffsplice. The DTU claim by limma is driven by the difference in mean transcript usage between cell types. Given the low abundance of the corresponding gene and the highly dispersed single-cell level observations, satuRn doesn’t identify the transcript as differentially used. **Panel B:** Transcript uniquely identified as differentially used by satuRn. Even though the mean difference in transcript usage between cell types is estimated to be 3%, satuRn claims significance given that the difference is stably supported by many cells with high gene-level expression levels. **Panel C:** Transcript uniquely identified as differentially used by limma diffsplice. The DTU claim by limma is driven by the raw mean difference in transcript usage between cell types. In contrast, satuRn takes into account that the Hsd11b1 Endou cells expressing the transcript at 0% usage have low gene-level count. The size of the dots (which represent individual cells) is weighted according to the total expression of the gene in that cell, i.e. the total gene count per cell. The yellow diamonds indicate the estimated mean usage of a transcript for each cell type, as estimated by satuRn. The cyan diamonds indicate the raw mean transcript usage levels per cell type.

Conversely, by looking at the transcripts that were highly ranked in our DTU list but lowly ranked in the top list of limma, we observe that our model is more likely to capture small changes in transcript usage that are stable across all cells and belong to genes that are highly expressed. An example of such a transcript is shown in Figure 8B. satuRn estimates a mean usage of 3% in Tnc cells and 6% in Hsd11b1 Endou cells. While this is only a minor change in transcript usage, satuRn still identifies this transcript as differentially used because the gene is highly expressed and the small change in usage is supported by a large number of cells. In case such small differences in usage are not considered biologically meaningful, it is possible to set a threshold on the minimal desired difference. Finally, by not taking into account gene abundances, limma is more influenced by outlying observations that have a low gene-level abundance (Figure 8C). Indeed, DTU claims by limma are driven by differences in raw mean usages of transcripts. In Figure 8C, the raw mean usage of the transcript is 77% in Tnc cells and 45% in Hsd11b1 Endou cells, as indicated by the cyan diamonds. By contrast, the mean usage estimate by satuRn, which takes into account that the Hsd11b1 Endou cells expressing the transcript at 0% usage have low gene-level count, is 83% for Tnc cells and 75% for Hsd11b1 Endou cells, as indicated by the gold diamonds.

We therefore argue that, given the above observations, the transcripts identified by satuRn should be considered more reliable, as they generally originate from genes containing more information for assessing DTU.

### Comparison to DoubleExpSeq

We additionally analyzed the dataset by Tasic *et al*. with DoubleExpSeq^20^. DoubleExpSeq identified a large number of DTU transcripts in all eight comparisons between cell types, ranging from 335 to 4580 DTU transcripts (Figure S12). This is consistent with our performance benchmarks, which already suggested that DoubleExpSeq becomes overly liberal in single-cell datasets with a large number of cells (Figures 4, S7, S8 and S9). We therefore expect many of these transcripts to correspond to false positives. Furthermore, this is reflected in the pathological distribution of p-values obtained by DoubleExpSeq, where p-values have a tendency to be small and therefore the analysis too liberal (Figure S15). Furthermore, as discussed in the benchmark studies, we could not adopt the empirical null strategy to improve the FDR control of DoubleExpSeq. Again, a large number of p-values equal 1 poses a problem for estimating the empirical null distribution (Figure S16).

While the results of DoubleExpSeq are likely to be overly liberal, the ranking of the transcripts (based on the p-values of the DTU analysis) might still be reasonable. In Figure 9, we observe a large overlap between the top 200 transcripts identified by satuRn in comparison #6 of the case study and the top 200 transcripts of DoubleExpSeq in that comparison. This overlap is considerably smaller with a limma diffsplice analysis.

**Figure 9:**
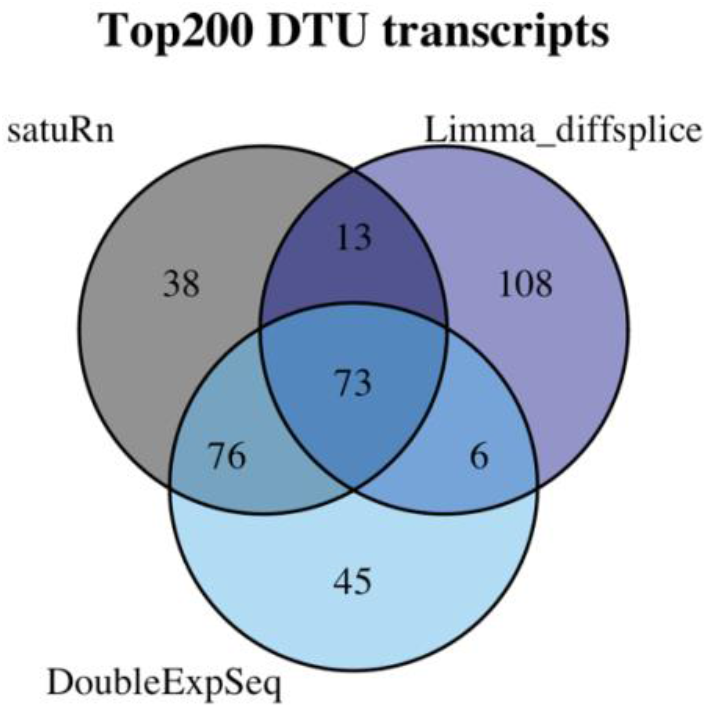
Venn diagram displaying the degree of overlap of the top 200 transcripts in comparison #6 of the case study in three DTU analysis tools. We observe that in the set of the top 200 transcripts identified by satuRn, 149 transcripts overlap with the top 200 list from DoubleExpSeq. In the top 200 list of limma diffsplice, 108 transcripts are present that were not in the top lists of satuRn or DoubleExpSeq.

Finally, we note that while DoubleExpSeq could still be used in this case study given the simple factorial design (using a single factor to assign each cell to a cell type), DoubleExpSeq cannot be used in multifactorial designs, for instance to compare expression levels across multiple cell types between multiple samples or treatment groups.

## Discussion

In this manuscript, we have proposed satuRn, a new software tool for DTU analysis. satuRn adopts a quasi-binomial GLM framework and obtains direct inference on DTU by modelling the relative usage of a transcript, in comparison to other transcripts from the same gene, between conditions of interest. We evaluated the performance of satuRn with respect to 7 other DTU methods on three simulated bulk RNA-seq datasets, a real bulk RNA-seq dataset and three real scRNA-seq datasets. These benchmarks underscored the strong performance of satuRn, as well as its ability to control the FDR close to the nominal level. In addition, we showed that satuRn scales seamlessly to the large data volumes that are produced in contemporary (sc-)RNA-seq experiments. Furthermore, given the underlying GLM framework, our method can handle complex experimental designs that are commonplace in scRNA-seq experiments. Finally, satuRn can extract biologically relevant information from a large scRNA-seq dataset that would have remained obscured in a canonical DGE analysis.

Since most sequencing reads map to multiple transcripts, quantification tools such as Salmon or kallisto only provide an estimate of the expected number of fragments originating from each transcript. Incorporating quantification uncertainty has recently been shown to improve results in differential expression analysis of single-cell RNA-seq datasets^44^. Currently, satuRn and all other DTU methods discussed in this manuscript, except for BANDITS^29^, neglect the uncertainty on this abundance estimate. BANDITS models the abundance uncertainty, however, it had a markedly lower performance than our method in our benchmark evaluation (Figure S1).

One challenge common to all DTU methods is that the power to detect differentially used transcripts depends strongly on the quality of the scRNA-seq dataset. This becomes clear when comparing the performances for the three different scRNA-seq benchmarks in this manuscript. The performances on the Darmanis^25^ dataset (Figure S9) are markedly lower than the performances on the other two datasets (Figures 4 and S8). A closer inspection of the Darmanis dataset showed that, after filtering, the transcript-level counts matrix contains a much larger percentage of zero counts than the other datasets. We also more frequently observed the scenario where the expression level of a gene could be attributed to a single isoform. This effectively causes the transcript usage to appear binary, with either 0% or 100% usages of a certain transcript. We argue that while this may reflect the true underlying biology, for instance through the process of transcriptional bursting^45,46^, it is more likely to be a technical artefact as a consequence of more shallow sequencing, given the lower percentage of binary usage profiles in the Chen and Tasic datasets. The supposedly binary expression of transcripts due to coverage-dependent bias and the use of more stringent filtering criteria to reduce this bias has already been comprehensively reported by Najar *et al*.^47^.

We conclude with the following recommendations for DTU analysis from an applied perspective. In case of small bulk RNA-seq datasets, satuRn, DEXSeq and DoubleExpSeq can be used interchangeably. In case of datasets with more complex designs that require the DTU model to incorporate additional covariates, e.g. batch effects, DoubleExpSeq cannot be used. For single-cell datasets, using DEXSeq will become infeasible in terms of scalability and DoubleExpSeq may give overly liberal results. As such, we recommend satuRn for performing DTU analyses in large bulk and single-cell RNA-seq datasets.

## Methods

### satuRn model

As input, satuRn requires a matrix of transcript-level expression counts, which may be obtained either through pseudo-alignment using, e.g.,, kallisto^1^, salmon^2^ or by classical alignment-based tools followed by transcript-level quantification (e.g. STAR^48,49^ and RSEM^50^). Let *Y*_*gti*_ denote the observed expression value for a given transcript *t = 1*, …, *T*_*g*_ of gene *g = 1*, …, *G* in cell or sample *i = 1*, …, *n*. The total expression of gene *g* in sample *i* can then be expressed as

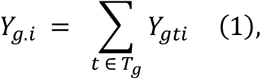

i.e. by taking the sum of expression values for all *T*_*g*_ transcripts belonging to gene *g* in sample *i*. The usage of transcript *t* in sample or cell *i* can then be estimated as

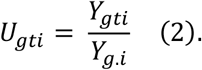

Next, we adopt a quasi-binomial (QB) generalized linear modelling (GLM) strategy to model DTU. As opposed to canonical maximum likelihood models, this quasi-likelihood modelling strategy only requires the specification of the first two moments of the response distribution, i.e. the mean and the variance. We define the mean of the QB model as

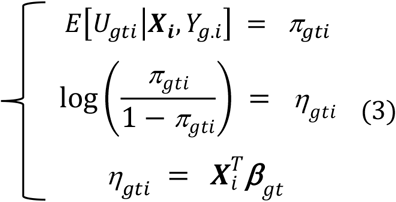

In this notation, *π*_*gti*_ is the expected probability of observing transcript *t* within the pool of transcripts (1, …, T_g_) belonging to gene *g* in sample *i* and, as such, corresponds to its expected usage for that sample. We model *π*_gti_ using a logit link *function*, where ***β***_t_ is a p x 1 column vector of regression parameters modeling the association between the average usage and the covariates for transcript *t*. Finally, 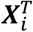 is a row in the n x p design matrix ***X*** that corresponds with the covariate pattern of sample *i*, with *p* the number of parameters of the mean model, i.e. the length of vector ***β***_t_.

The variance of the QB model can be described as

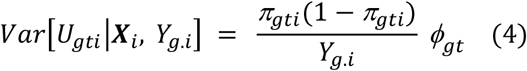

with *Y*_*g*.*i*_*π*_*gti*_(1 − *π*_*gti*_) the canonical variance of the binomial distribution and *ϕ*_gt_ a transcript-specific overdispersion parameter to describe additional variance in the data with respect to the binomial variance. We adopt the empirical Bayes procedure from Smyth *et al*.^23^, as implemented in the *squeezeVar* function of the *limma* Bioconductor R package, to stabilize the estimates of *ϕ*_gt_ by borrowing information across transcripts, which is adopted in the default edgeR quasi-likelihood workflow for bulk RNA-seq data^22^ Note that stabilizing the dispersion estimation is particularly useful in datasets with a small sample size.

Taken together, the quasi-binomial thus allows us to model the log-odds of drawing a particular transcript *t* from the pool of transcripts in the corresponding gene *g* across samples. The intercept also has an interpretation of a log-odds and the remaining mean model parameters are log-odds ratios, which may thus be interpreted in terms of differential transcript usage. We adopt t-tests that are computed based on the log-odds ratio estimates of the QB model and the posterior variance, as obtained from the empirical Bayes procedure. P-values are computed assuming a t-distribution under the null hypothesis with posterior degrees of freedom calculated as the sum of the residual degrees of freedom and the prior degrees of freedom from the empirical Bayes procedure.

For bulk analyses, the implementation of satuRn as described above provides a high performance and a good control of the FDR. However, for single-cell datasets we observed that our inference is too liberal (Figure S10), which could suggest that the theoretical null, the t-distribution, is no longer valid. Indeed, in large-scale inference settings, failure of the theoretical null distribution is often observed. Efron^52^ (Chapter 6) describes four reasons why the theoretical null distribution may fail; failed mathematical assumptions, correlation across features (transcript expression), correlation across subjects (samples or cells), and unobserved confounders in observational studies. To avoid these issues, Efron proposes to exploit the massive parallel data structure of omics datasets to empirically estimate the null distribution of the test statistics^53^. To this end, Efron converts the test statistic to z-scores, which should follow a standard normal distribution under the theoretical null, and then proposes to approximate the empirical null distribution with a normal distribution with unknown mean (*μ*^*^) and standard deviation (*σ*^*^), which can be estimated by maximum likelihood on a subset of the test statistics near zero.

As such, we first convert the two-sided p-values to z-scores according to

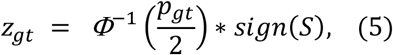

with Ф the cumulative distribution function for the standard normal distribution, *p*_*gt*_ the original two-sided p-value indicating the statistical significance of differential usage of transcript *t* from gene *g* between the conditions of interest, sign(*S*) the sign of the t-test statistic S and *z*_*gt*_ the resulting z-score. Next, we adopt the maximum likelihood procedure, implemented in the *locfdr* function of the locfdr R package from CRAN^54^, to estimate the mean *μ*^*^ and standard deviation *σ*^*^of the empirical null distribution. Based on these estimates, we recompute the z-scores and corresponding p-values as follows

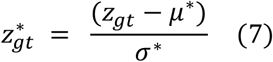

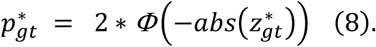

Finally, the resulting (empirical) p-values are corrected for multiple testing with the FDR method of Benjamini and Hochberg^30^. As opposed to the original p-values that were calculated based on the theoretical null distribution for the t-statistics, we found that this procedure allows for a better FDR control in single-cell applications.

### DTU tools literature

Below we provide a brief description of each of the DTU methods from the literature that were included in the performance benchmarks of this paper. For more details, we refer to the respective original publications. Note that all methods were run with the current default settings.

### DEXSeq

DEXSeq^19^ (R package version 1.32.0) takes as input a transcript-level expression matrix *Y*_*ti*_, with *T* transcripts (rows) and *n* samples or cells (columns). Next, a matrix of complementary counts *C*_*ti*_ is calculated, which defines how many reads map to any of the other transcripts of the same gene as respective transcript *t* in cell *i*. DEXSeq then augments the original expression matrix *Y*_*ti*_ by concatenating it with the complementary counts *C*_*ti*_, hence doubling the number of columns of the original count matrix. A negative binomial generalized linear model (GLM) is fitted to each transcript in the augmented count matrix as follows

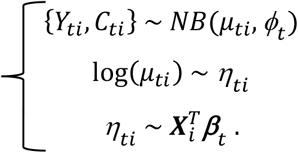

In the specification of the GLM, 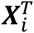 corresponds to row i of design matrix **X**, which defines a covariate pattern that (i) links the transcript-level count matrix to the complementary counts through sample-level intercepts, and (ii) specifies the design of the experiment. Inference on DTU is obtained by testing an interaction effect that assesses if the log fold change between transcript *t* and all other transcripts in its corresponding gene changes between the conditions of interest (e.g. treatment) with a likelihood ratio test. It is important to note that the estimation of sample-level intercepts is required because of the concatenation of the two count matrices. As a consequence, DEXSeq scales quadratically with the number of samples or cells in the data. The lack of scalability is thus inherent to the parametrization of DEXSeq, putting a severe burden on the utility of DEXSeq for DTU analysis in large datasets, as displayed in Figure 1.

### DoubleExpSeq

DoubleExpSeq^20^ (R package version 1.1) assumes a double binomial distribution for each transcript. The double binomial distribution is a member of the double exponential family of distributions described by Efron^55^, which are extensions of one-parameter exponential family distributions that allow for a more flexible variance structure through introduction of an additional dispersion parameter. DoubleExpSeq adopts a bespoke empirical Bayes procedure for computing shrinkage estimates of the dispersion parameter of the double binomial distribution. The double binomial models the log-odds of drawing a particular transcript *t* from the pool of transcripts in the corresponding gene *g* across samples. The intercept thus has an interpretation of a log-odds and the remaining mean model parameter(s) are log-odds ratios, which may thus be interpreted in terms of differential transcript usage. The significance of the mean model parameter(s) are tested using a likelihood ratio test. Importantly, the current implementation of DoubleExpSeq does not allow for modeling multifactorial designs and cannot make use of parallel computing.

### DRIMSeq

DRIMSeq^21^ (R package version 1.14.0) assumes that the transcript-level expression counts marginally follow a Dirichlet multinomial distribution (DM), where the Dirichlet conjugate prior is used to account for overdispersion with respect to the multinomial distribution. The most important consequence of treating transcript expression as a realization of a multinomial distribution, is that the correlations between expression of transcripts derived from the same gene are directly accounted for. In the DRIMSeq framework, the total count for a gene is considered fixed, and the quantity of interest is the change in proportion of each transcript within a gene between groups of samples or cells. More specifically, DRIMSeq uses a likelihood ratio test to determine if the transcript ratios of a gene, which are modelled by the multinomial, are different between conditions of interest.

### Limma diffsplice

Limma diffsplice (limma, R package version 3.42.2) is a built-in functionality described in the current user’s guide of the limma Bioconductor R package^23^. Limma was originally devised for analyzing microarray data but can also be used for RNA-Seq data with the limma-voom method^56^. Limma-voom fits a linear model to the log-transformed (normalized) transcript-level count matrix, while adjusting for heteroskedasticity via weighted regression, where the observation weights are computed from the observed variance-mean relationship. Limma diffsplice then uses a series of t-tests to assess DTU at the transcript level by comparing the log-fold change in expression of transcript *t* with the average log-fold change in the expression of all transcripts belonging to the same gene as transcript *t*.

### EdgeR diffsplice

EdgeR diffsplice (edgeR, R package version 3.28.1) is a built-in functionality described in the vignettes of the edgeR Bioconductor R package, which was last revisited by Chen et al.^22^. The edgeR diffsplice function fits a negative binomial GLM for each transcript and tests for differential transcript usage by comparing the obtained log-fold changes for each respective transcript within a gene with the log-fold change of the entire gene. If the log-fold change for a certain transcript is significantly different from those of the other transcripts in the gene, it is flagged as differentially used. Note that the negative binomial GLMs can be fit using a canonical likelihood-based approach or using a quasi-likelihood. We adopted the likelihood-based approach as it consistently displayed higher performances (data not shown). In this setting, inference is obtained using a likelihood ratio test.

### NBSplice

NBSplice^24^ (R package version 1.4.0) fits a negative binomial GLM for each transcript in the dataset. In contrast to e.g. DEXSeq, the mean transcript-level expression (i.e. the mean parameter of the negative binomial model) is taken as the product of the mean gene-level expression value and the observed percentual usage of the transcripts within its corresponding gene. The GLM framework of NBSplice is structured such that DTU between groups of interest can be tested using a likelihood ratio test, where the full model contains an isoform-condition interaction term that is omitted in the null model. Note that in our benchmarks the NB GLM estimation procedure of NBSplice fails to converge when there is a large fraction of zero counts in the data. As a consequence, NBSplice was omitted from the performance benchmarks on single-cell data and from the scalability benchmarks, as the latter also make use of single-cell data.

### BANDITS

BANDITS^29^ (R package version 1.3.2) adopts a Bayesian hierarchical model with a Dirichlet-multinomial to explicitly model the sample-to-sample variability between biological replicates. In addition to the transcript-level count matrix, equivalence class counts are used as input to the BANDITS algorithm. As described by Bray et al.^1^, an equivalence class for a (transcriptomics) read is a multi-set of transcripts associated with that read. As such, an equivalence class represents the transcripts from which a read could have originated. BANDITS leverages the information conveyed by the equivalence class counts to model the uncertainty arising from reads mapping to multiple transcripts. In brief, the allocation of reads to transcripts is treated as a latent variable that is sampled jointly with the parameters of the Dirichlet-multinomial; sampling of these parameters is done with a Markov chain Monte Carlo algorithm. As such, BANDITS allows for modeling the mean relative usage of each transcript within its corresponding gene across samples/cells, while accounting for quantification uncertainty. In addition, BANDITS also accounts for differences in transcript length. Finally, BANDITS tests for DTU (at the transcript level) by performing univariate Wald tests.

### Filtering

We adopted two different strategies for filtering transcripts in each of the RNA-seq datasets in the performance benchmarks.

The first filtering strategy uses the *filterByExpr* function implemented in edgeR^57^. This filtering strategy only retains transcripts that have at least an expression level of *min*.*count* counts-per-million (CPM, calculated as the number of read counts divided by the total number of reads in the dataset and multiplied by one million) in at least *n* samples or cells. In addition, the sum of the CPM of the transcript across all cells or samples must be at least *min*.*total*.*count*. For the bulk RNA-seq datasets, we use the default settings (*min*.*count* = 10, n = min(10, 0.7*sample size of the smallest group in the comparison) and *min*.*total*.*count* = 10). For the scRNA-seq datasets, the settings are adjusted to; *min*.*count* = 1 (as requiring a transcript to be expressed in all single-cells is a stringent criterium), n = 0.5*sample size of the smallest group in the comparison and *min*.*total*.*count* = 0. In addition, if only one transcript of a gene passes this filtering criterion, it is omitted from the analysis, as DTU analysis is meaningless when only one transcript is retained. As such, we specifically set the parameters to generate a very lenient filtering criterium.

The second filtering strategy uses the *dmFilter* function implemented in DRIMSeq^21^. This filter is more stringent and specifically designed for DTU analysis. The filtering process can be thought of as proceeding in three steps. Let *n*_*s*_ be the number of samples or cells in the smallest group. The first step requires the transcripts to have a count of at least 10 in at least *n*_*s*_ samples. The second filtering step requires the transcript to make up at least 10% of the total count of its corresponding gene in at least *n*_*s*_ samples. The third filtering step removes all transcripts for which the corresponding gene has a count below 10 in any of the samples or cells in the dataset. Again, if only one transcript of a gene passes this filtering criterion, it is omitted from the analysis.

### Bulk simulation study

To evaluate the performance of the different DTU analysis methods, we first adopt three simulated bulk RNA-seq datasets from previous publications: the simulated dataset from Love et al.^18^ (dataset 1) and both the Drosophila melanogaster (dataset 2) and Homo sapiens (dataset 3) simulation studies from Van den Berge et al.^31^. All three datasets were generated based on parameter values obtained from real RNA-seq samples, to mimic real RNA-seq data as close as possible.

Notably, there is a subtle difference in how DTU is introduced between the two simulation frameworks. For dataset 1, the origin of DTU is twofold: On the one hand, DTU was specifically introduced by swapping the transcript-per-million (TPM) abundances between two expressed isoforms. On the other hand, DTU was also obtained as a consequence of introducing DTE, where a single expressed isoform was induced to be differentially expressed at a certain log fold change, which leads to DTU if this transcript belongs to a gene expressing multiple isoforms. For datasets 2 and 3, there is only one source of DTU. The number of differentially used transcripts within a gene was sampled ranging from a minimum of 2 up to a random number drawn from a binomial distribution with size equal to the number of transcripts and success probability 1/3. DTU was introduced by swapping the TPM abundances between the differentially used transcripts. As such, the latter framework allows for differential usage of multiple transcripts of the same gene, which is not possible with the framework used for generating dataset 1. Additionally, dataset 1 uses salmon^2^ (version 1.1.0) for estimating transcript-level abundances, whereas datasets 2 and 3 were quantified with kallisto^1^ (version 0.46.2).

### Real bulk study

We evaluate the performance of the different DTU methods on real bulk RNA-seq data, by subsampling a homogeneous set of samples from the large bulk RNA-seq dataset available from the Genotype-Tissue Expression (GTEx) consortium^34^ release version 8. Nine datasets were generated non-parametrically. More specifically, we first selected samples from adrenal gland tissue that were extracted with the RNA extraction method “RNA Extraction from Paxgene-derived Lysate Plate Based”. From the remaining samples we subsampled 9 datasets, comprising 3 repeats for each of 3 sample sizes; *5 versus 5, 20 versus 20* and *50 versus 50* samples. Next, DTU is artificially introduced with the swapping strategy that is described in the bulk simulation study paragraph of the Methods section of this paper. The GTEx data was quantified with RSEM^50^ version 1.3.0.

### Real single-cell study

We evaluate the performance of the different DTU methods on real scRNA-seq datasets. These scRNA-seq datasets were generated non-parametrically by subsampling a homogeneous set of cells from three real scRNA-seq datasets^25,28,35^, after which DTU is artificially introduced by the swapping strategy that is described in the bulk simulation study paragraph of the Methods section of this paper.

For the dataset of Chen *et al*.^28^, which was used to construct Figures 4 and S7, we selected a homogeneous population of cells by considering only the EpiStem cells of female mice, resulting in a dataset of 120 cells. From this homogeneous population of cells, we then subsampled 6 datasets, comprising 3 repeats for each of 2 sample sizes: *20 versus 20* and *50 versus 50* cells. Next, DTU was artificially introduced with the swapping strategy that is described in the bulk simulation study paragraph of the Methods section of this paper. Finally, we adopted either edgeR or DRIMSeq for filtering.

The other two scRNA-seq datasets were generated analogously. For the dataset of Tasic *et al*.^35^, which was used to construct Figure S8 in the main manuscript, we selected a homogeneous population of cells by considering only the Lamp5 cells in the anterior lateral motor cortex of mice without any eye conditions, resulting in a dataset of 897 cells. After introducing DTU, we randomly subsampled 20, 75 or 200 cells from each group. For the dataset of Darmanis *et al*.^25^, which was used to construct Figure S9, we selected the immune cells that clustered together in tSNE cluster 8 of the original publication, resulting in a dataset of 248 cells. After introducing DTU, we randomly subsampled 20, 50 or 100 cells from each group.

### Case study DGE analysis

We perform a DGE analysis on a subset of the Tasic single-cell dataset^35^, i.e. between different the cell types originating from the ALM and VISp regions of the glutamatergic L5 IT subclass. We use the quasi-likelihood method of edgeR^32^ to model the gene expression profiles and additionally adopt the edgeR *glmTreat* function to test differential expression against a log2-fold change threshold (log2-fold change = 1). Statistical significance was evaluated at the 5% FDR level.

### Performance assessment

We assess the performance of different DTU methods on a bulk simulation dataset with scatterplots of the true positive rate (TPR) versus the false discovery rate (FDR), according to the following definitions:

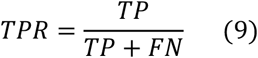

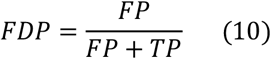

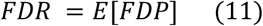

where FN, FP and TP denote the numbers of false negatives, false positives and true positives, respectively. The FDR-TPR curves are constructed using the Bioconductor R package ICOBRA^58^.

### Scalability benchmark

The scalability benchmark was run on subsets of the Chen scRNA-seq dataset^28^, which contains 617 cells in total. For the scalability benchmark with respect to the number of cells in the dataset, we randomly subsample a certain number of cells (8, 16, 32, 64, 128 or 256 cells per group) from the dataset (without introducing DTU or selecting specific homogeneous cell populations). Next, we filter this subsample using the edgeR-based filtering criterion. This was done to remove very lowly abundant transcripts, which may otherwise cause problems in the parameter estimation procedure. From the remaining transcripts, we randomly subsampled to a total of 30.000 transcripts before running the DTU analysis. To allow for a scalability benchmark of BANDITS, which scales poorly to the number of transcripts (Figure 5B), we also generated a dataset with only 1.000 transcripts (Figure S1).

For the scalability benchmark with respect to the number of transcripts, we randomly sampled two groups of 16 cells from the dataset. After applying the edgeR-based filter, we sampled 8 distinct numbers of transcripts: 1.000, 2.000, 5.000, 10.000, 15.000, 20.000, 25.000, 30.000 and 35.000 prior to the DTU analysis.

All scalability benchmarks were run on a single core of a virtual machine with an Intel(R) Xeon(R) CPU E5-2420 v2 (2.20GHz, Speed: 2200 MHz) processor and 30GB RAM.

## Acknowledgements

The authors would like to thank Milan Malfait for his suggestions and comments throughout this project.

## Data and code availability

satuRn is implemented in an R package that is available at https://github.com/statOmics/satuRn and will be submitted to the Bioconductor project. All the scripts that are required to reproduce the analyses and figures that are used for this publication can be retrieved from https://github.com/statOmics/satuRnPaper. On this GitHub page, a Zenodo link will provided from which the raw data and intermediate results of our analyses can be downloaded.

## Supplementary Figures

**Figure S1:**
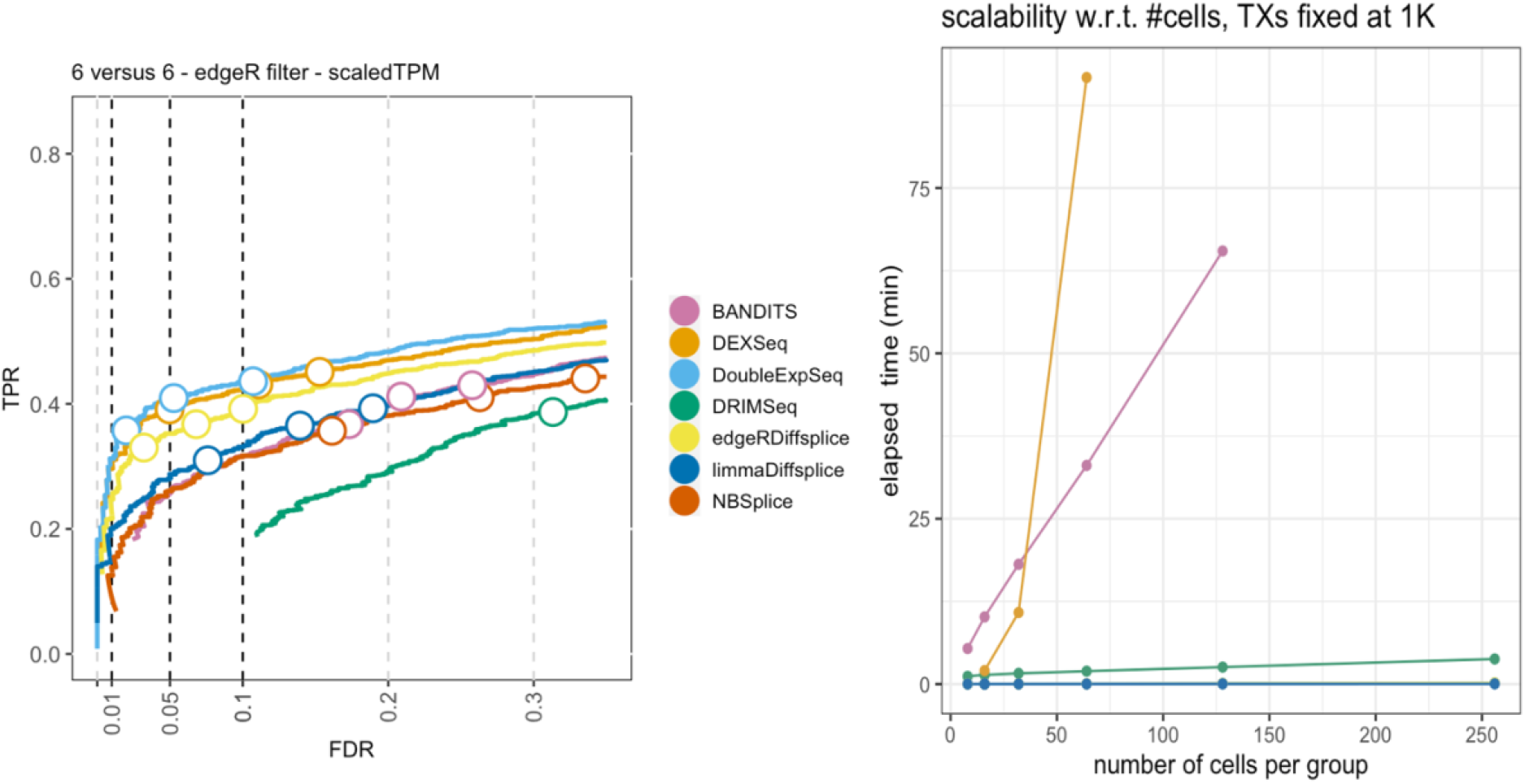
Performance and scalability evaluation on a subset of the Love et al. dataset. To allow for a performance and scalability evaluation of BANDITS, which does not scale to datasets with a large number of transcripts, we here perform a DTU analysis for the *6 versus 6* samples dataset of Love et al. with only 1000 transcripts. **Left panel: performance evaluation**. The results are in line with those of Figure 1A. The performance of BANDITS is indicated in pink. **Right panel: Scalability evaluation**. BANDITS scales linearly with respect to the number of cells (or samples) in the dataset. The slope of the linear trend, however, is considerably larger than those of the other DTU methods that scale linearly. Note that the profiles of limma diffsplice, edgeR diffsplice and DoubleExpSeq overlap in this figure.

**Figure S2:**
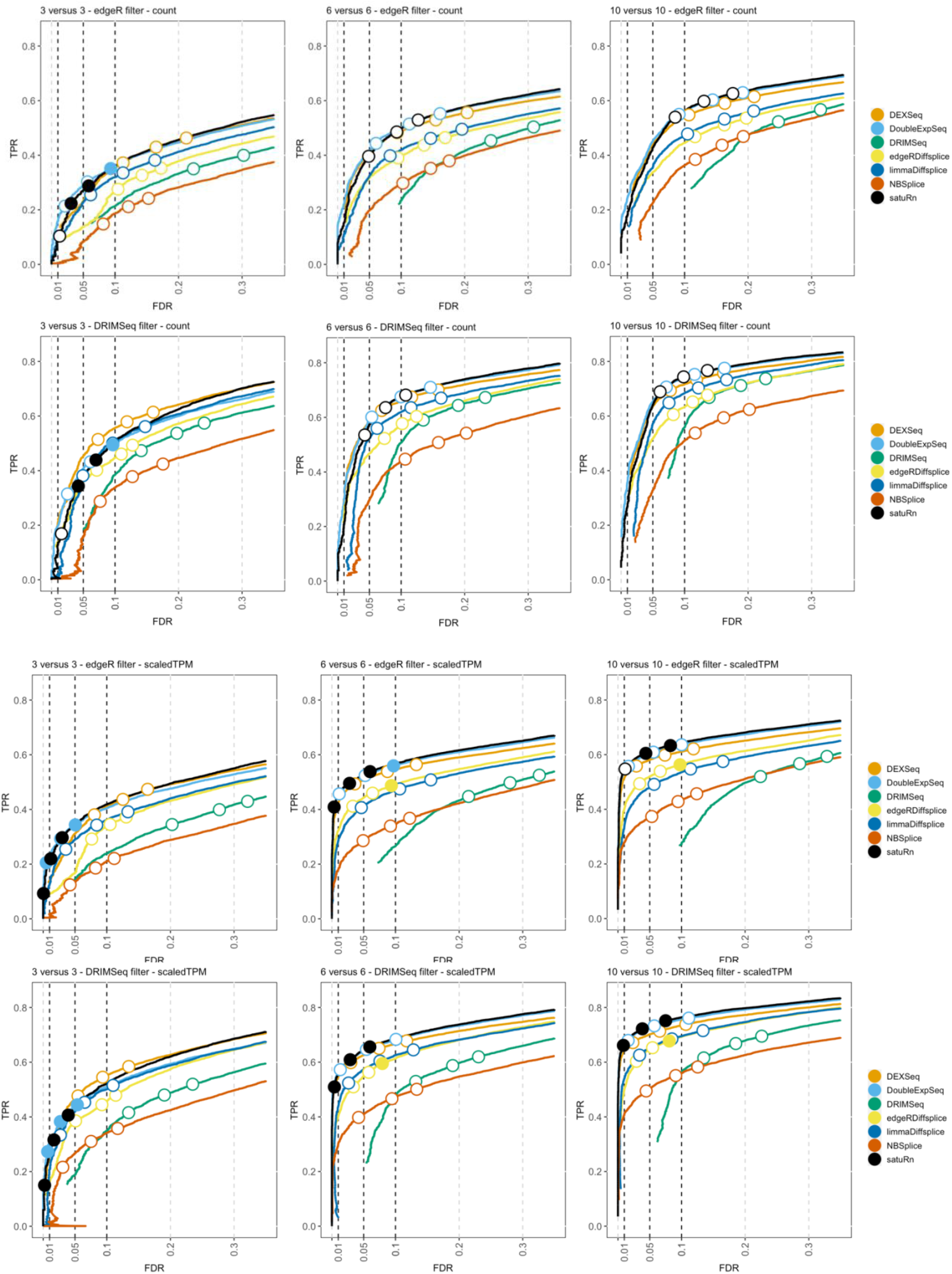
Performance evaluation of satuRn on different subsamples of the simulated bulk RNA-Seq dataset by Love et al. FDR-TPR curves visualize the performance of each method by displaying the sensitivity of the method (TPR) with respect to the false discovery rate (FDR). The three circles on each curve represent working points when the FDR level is set at nominal levels of 1%, 5% and 10%, respectively. The circles are filled if the empirical FDR is equal or below the imposed FDR threshold. We subsampled two-group comparisons according to three different samples sizes; a *3 versus 3, 6 versus 6* and *10 versus 10* comparison, as denoted in the panel titles. The benchmark was performed both on the raw counts **(rows 1 and 2)** or on scaled transcripts-per-million (TPM) **(rows 3 and 4)** as imported with the Bioconductor R package tximport^59^. We additionally adopted two different filtering strategies: an edgeR-based filtering **(rows 1 and 3)** and a DRIMSeq-based filtering **(rows 2 and 4)**. Overall, the performance of satuRn is on par with those of the best tools in the literature, DEXSeq and DoubleExpSeq. In addition, satuRn achieves a better control of the FDR on all datasets. For extremely small sample size, i.e. the *3 versus 3* comparison, the performance is slightly below that of DEXSeq, and inference does become slightly too conservative. Note that, as expected, the performances increase with increasing sample size, and a higher performance is achieved with the more stringent DRIMSeq filtering criterion (see Methods), which goes at the cost of retaining fewer transcripts for DTU analysis. Finally, we note that the performances and FDR control are consistently higher for the scaled TPM data as compared to the raw counts. Note that this was only observed for this particular dataset.

**Figure S3:**
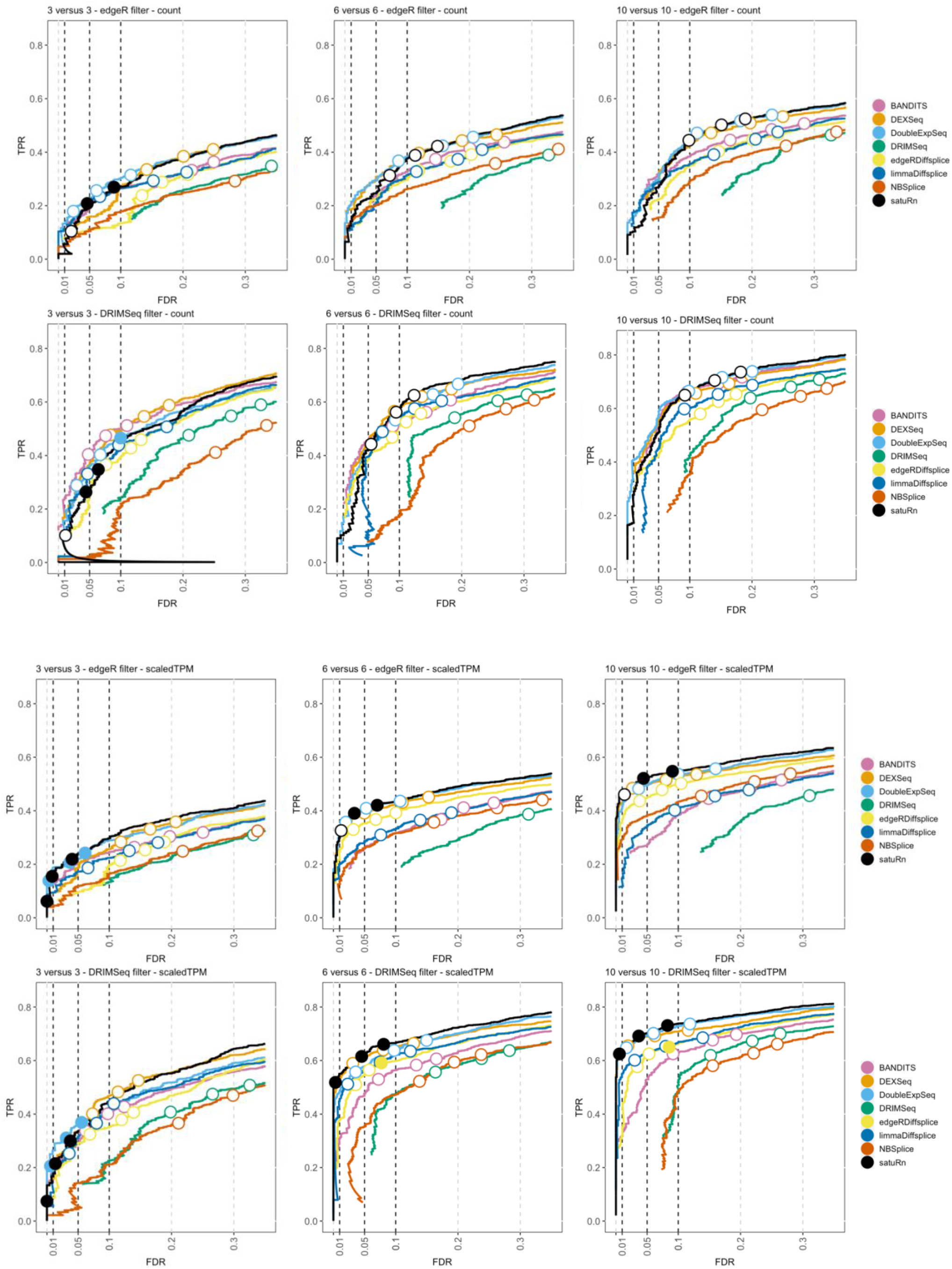
Performance evaluation on different subsamples of the simulated bulk RNA-Seq dataset by Love et al. with a reduced number of transcripts to allow for a comparison with BANDITS. FDR-TPR curves visualize the performance of each method by displaying the sensitivity of the method (TPR) with respect to the false discovery rate (FDR). The three circles on each curve represent working points when the FDR level is set at nominal levels of 1%, 5% and 10%, respectively. The circles are filled if the empirical FDR is equal or below the imposed FDR threshold. We subsampled two-group comparisons according to three different samples sizes; a *3 versus 3, 6 versus 6* and *10 versus 10* comparison, as denoted on top of the panels. The benchmark was performed both on the raw counts **(rows 1 and 2)** or on scaled transcripts-per-million (TPM) **(rows 3 and 4)** as imported with the Bioconductor R package tximport^59^. We additionally adopted two different filtering strategies: an edgeR-based filtering **(rows 1 and 3)** and a DRIMSeq-based filtering **(rows 2 and 4)**. Note that, in contrast to Figure S2, we additionally randomly subsampled 1000 genes (∼3000-5000 transcripts) after filtering, in order to reduce the number of transcripts in the data and thereby allowing for a DTU analysis with BANDITS. In concordance with Figure S2, the performance of satuRn is on par with the best tools of the literature with a better control of the FDR in general. While the performance of BANDITS is good for the settings for which it was originally developed, (i.e., small datasets with a stringent filtering criterium), its performance is reduced in larger, more leniently filtered datasets and inference is also overly liberal in these settings. In addition, while all other methods perform much better on the scaledTPM data (rows 3 and 4) than on the raw count data (rows 1 and 2), BANDITS has a similar performance on both input data types. This can be explained by the fact that BANDITS inherently corrects for differences in transcript length, even when raw counts are used as an input.

**Figure S4:**
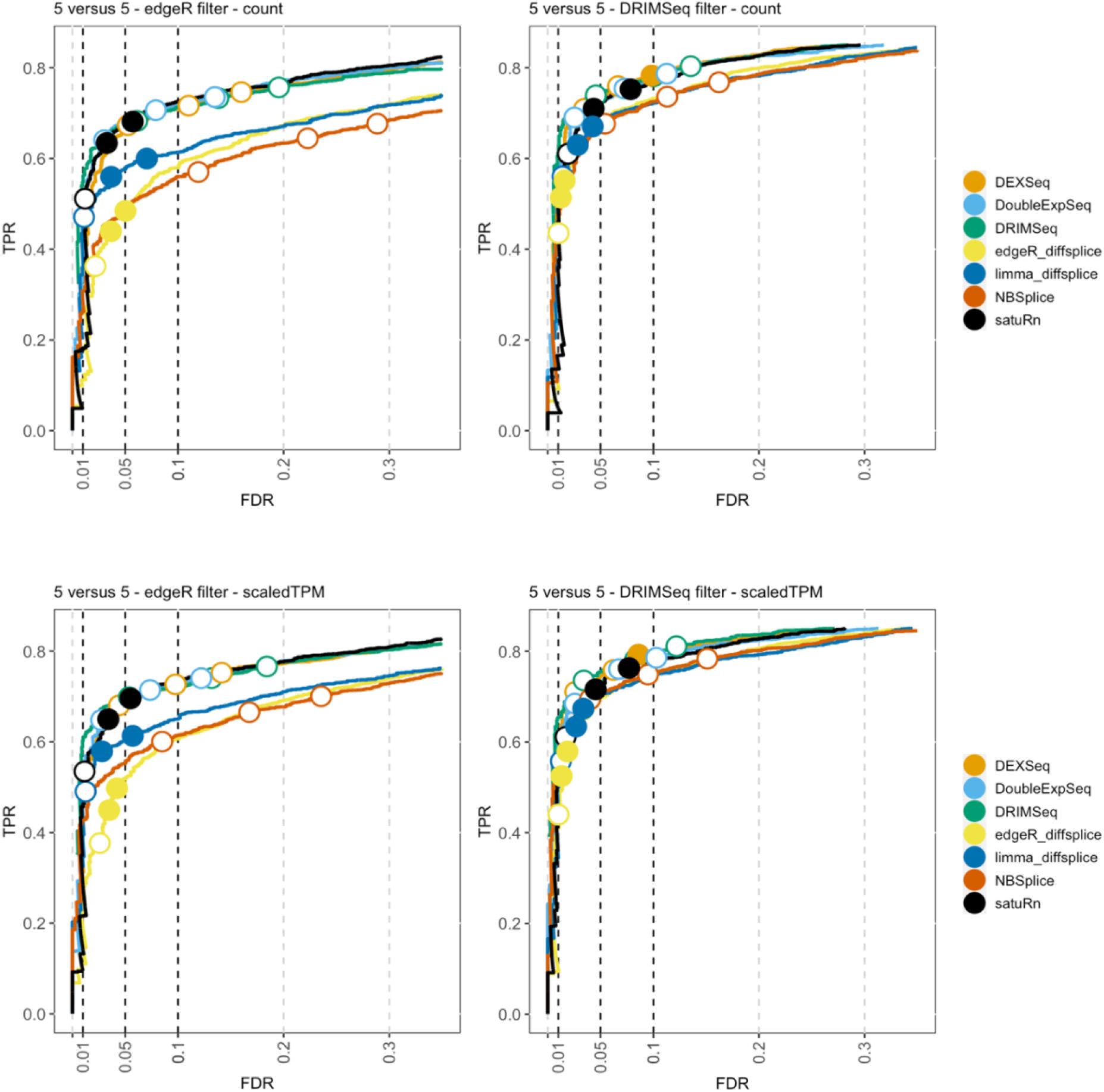
Performance evaluation of satuRn on the “Dmelanogaster” simulated bulk RNA-Seq dataset by Van den Berge et al. FDR-TPR curves visualize the performance of each method by displaying the sensitivity of the method (TPR) with respect to the false discovery rate (FDR). The three circles on each curve represent working points when the FDR level is set at nominal levels of 1%, 5% and 10%, respectively. The circles are filled if the empirical FDR is equal or below the imposed FDR threshold. The benchmark was performed both on the raw counts **(row 1)** and on scaled TPM **(row 2)** as imported with the Bioconductor R package tximport^59^. We additionally adopted two different filtering strategies; an edgeR-based filtering **(column 1)** and a DRIMSeq-based filtering **(column 2)**. Overall, the performance of satuRn is on par with those of the best tools in the literature, DEXSeq and DoubleExpSeq. In contrast to the performance evaluation on the dataset by Love et al. (Figures 1A and S2), there is a limited difference in performances based on the data input type (i.e., counts versus scaled TPM), and DRIMSeq also performs well on these datasets.

**Figure S5:**
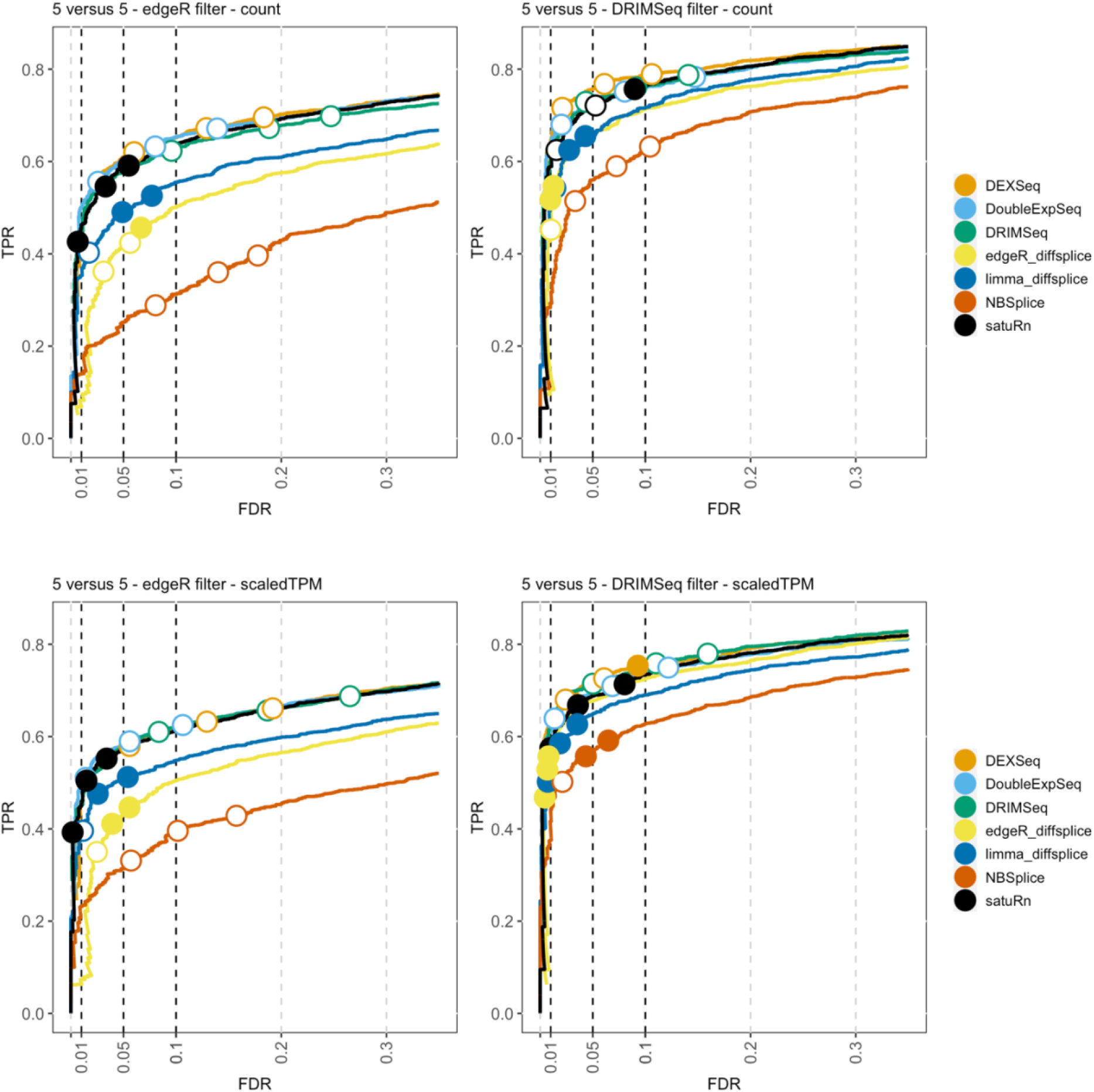
Performance evaluation of satuRn on the “Hsapiens” simulated bulk RNA-Seq dataset by Van den Berge et al. FDR-TPR curves visualize the performance of each method by displaying the sensitivity of the method (TPR) with respect to the false discovery rate (FDR). The three circles on each curve represent working points when the FDR level is set at nominal levels of 1%, 5% and 10%, respectively. The circles are filled if the empirical FDR is equal or below the imposed FDR threshold. The benchmark was performed both on the raw counts **(row 1)** and on scaled TPM **(row 2)** as imported with the Bioconductor R package tximport^59^. We additionally adopted two different filtering strategies; an edgeR-based filtering **(column 1)** and a DRIMSeq-based filtering **(column 2)**. Overall, the performance of satuRn is on par with those of the best tools in the literature, DEXSeq and DoubleExpSeq. In contrast to the performance evaluation on the dataset by Love et al. (Figures 1A and S2),), there is a limited difference in performances based on the data input type (i.e., counts versus scaled TPM), and DRIMSeq also performs well on these datasets.

**Figure S6:**
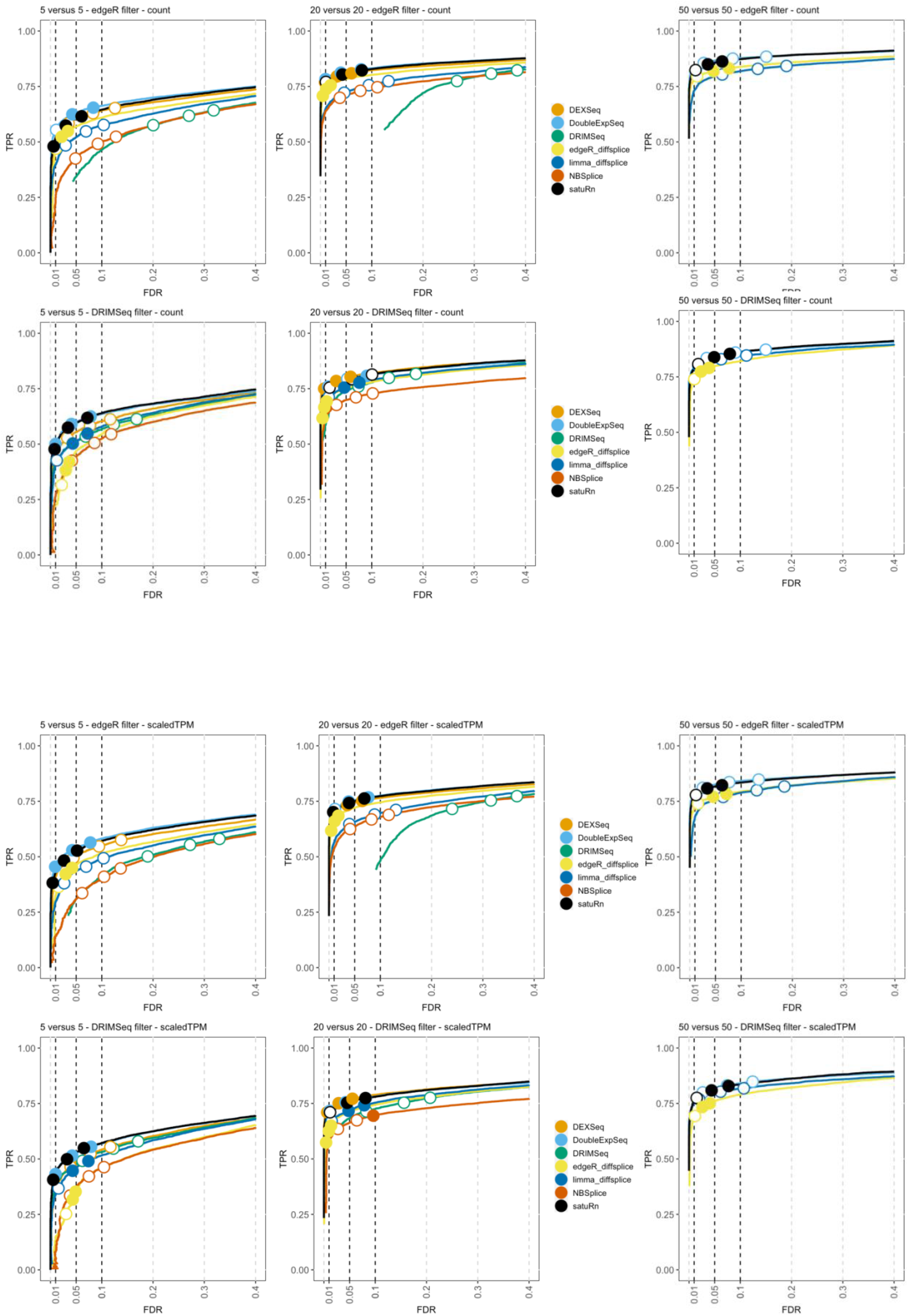
Performance evaluation of satuRn on the GTEx bulk RNA-Seq dataset. FDR-TPR curves visualize the performance of each method by displaying the sensitivity (TPR) with respect to the false discovery rate (FDR). The three circles on each curve represent working points when the FDR level is set at nominal levels of 1%, 5% and 10%, respectively. The circles are filled if the empirical FDR is equal or below the imposed FDR threshold. The benchmark was performed both on the raw counts **(rows 1 and 2)** or on scaled transcripts-per-million (TPM) **(rows 3 and 4)** as imported with the Bioconductor R package tximport^59^. We additionally adopted two different filtering strategies; an edgeR-based filtering **(rows 1 and 3)** and a DRIMSeq-based filtering **(rows 2 and 4)**. The performance of satuRn is on par with the best tools from the literature, DEXSeq and DoubleExpSeq. In addition, satuRn consistently provides a stringent control of the FDR, while DoubleExpSeq becomes more liberal with increasing sample sizes. Note that DEXSeq, DRIMSeq and NBSplice were omitted from the largest comparison, as these methods do not scale to large datasets (Figure1).

**Figure S7:**
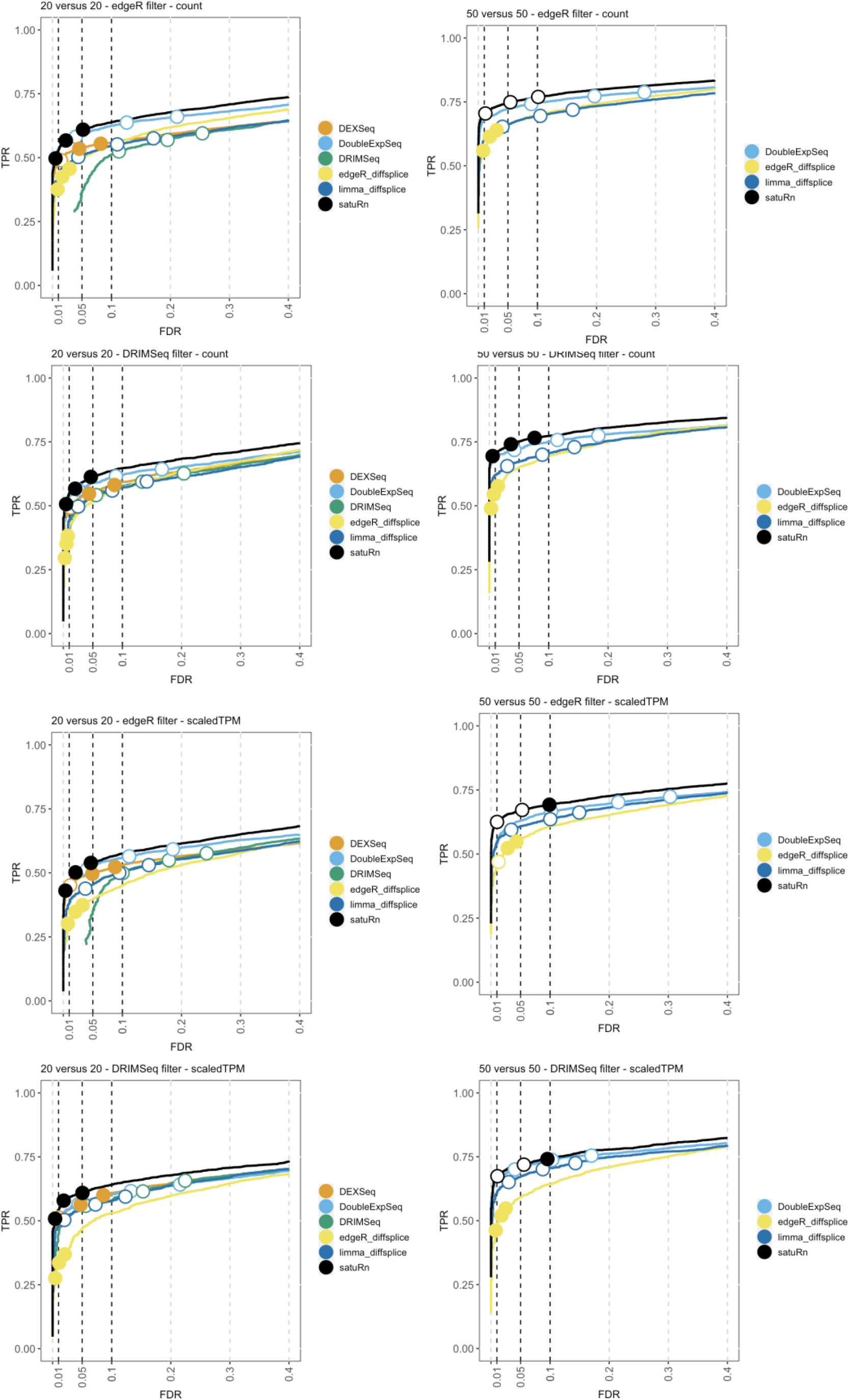
Performance evaluation of satuRn on the real scRNA-Seq dataset by Chen et al. FDR-TPR curves visualize the performance of each method by displaying the sensitivity of the method (TPR) with respect to the false discovery rate (FDR). The three circles on each curve represent working points when the FDR level is set at nominal levels of 1%, 5% and 10%, respectively. The circles are filled if the empirical FDR is equal or below the imposed FDR threshold. The benchmark was performed both on the raw counts **(rows 1 and 2)** or on scaled transcripts-per-million (TPM) **(rows 3 and 4)** as imported with the Bioconductor R package tximport^59^. We additionally adopted two different filtering strategies; an edgeR-based filtering **(rows 1 and 3)** and a DRIMSeq-based filtering **(rows 2 and 4)**. The performance of satuRn is at least on par with the best tools from the literature. Note that the performance of DEXSeq is clearly lower. In addition, our method consistently controls the FDR close to its imposed nominal FDR threshold, while DoubleExpSeq becomes more liberal with increasing sample sizes. DEXSeq and DRIMSeq were omitted from the largest comparison (two groups with 50 cells each), as these methods do not scale to large datasets (Figure 1). NBSplice was omitted from all comparisons, as it does not converge on datasets with many zeros, such as scRNA-Seq datasets.

**Figure S8:**
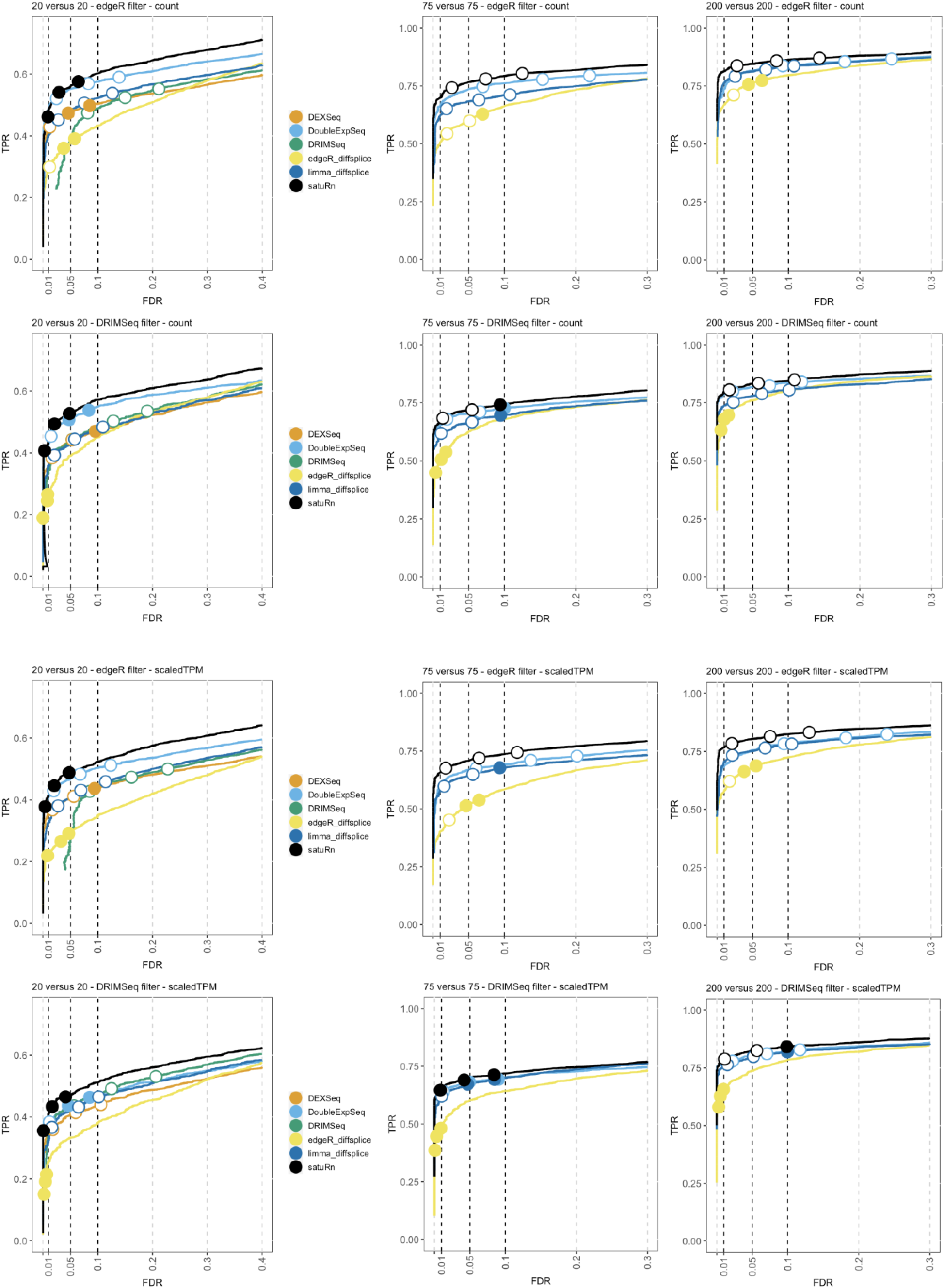
Performance evaluation of satuRn on the real scRNA-Seq dataset by Tasic et al. FDR-TPR curves visualize the performance of each method by displaying the sensitivity of the method (TPR) with respect to the false discovery rate (FDR). The three circles on each curve represent working points when the FDR level is set at nominal levels of 1%, 5% and 10%, respectively. The circles are filled if the empirical FDR is equal or below the imposed FDR threshold. We generated three two-group comparisons of 20, 75 and 200 cells each (left, middle and right panel, respectively). The benchmark was performed both on the raw counts **(rows 1 and 2)** or on scaled transcripts-per-million (TPM) **(rows 3 and 4)** as imported with the Bioconductor R package tximport^59^. We additionally adopted two different filtering strategies; an edgeR-based filtering **(rows 1 and 3)** and a DRIMSeq-based filtering **(rows 2 and 4)**. Overall, satuRn slightly outperforms DoubleExpSeq, the best tools from the literature. Note that the performance of DEXSeq is clearly lower. In addition, our method consistently controls the FDR close to its imposed nominal FDR threshold, while DoubleExpSeq becomes more liberal with increasing sample sizes. DEXSeq and DRIMSeq were omitted from the largest comparison (two groups with 75 cells and 200 cells each, respectively), as these methods do not scale to large datasets (Figure 1). NBSplice was omitted from all comparisons, as it does not converge on datasets with many zeros, such as scRNA-Seq datasets.

**Figure S9:**
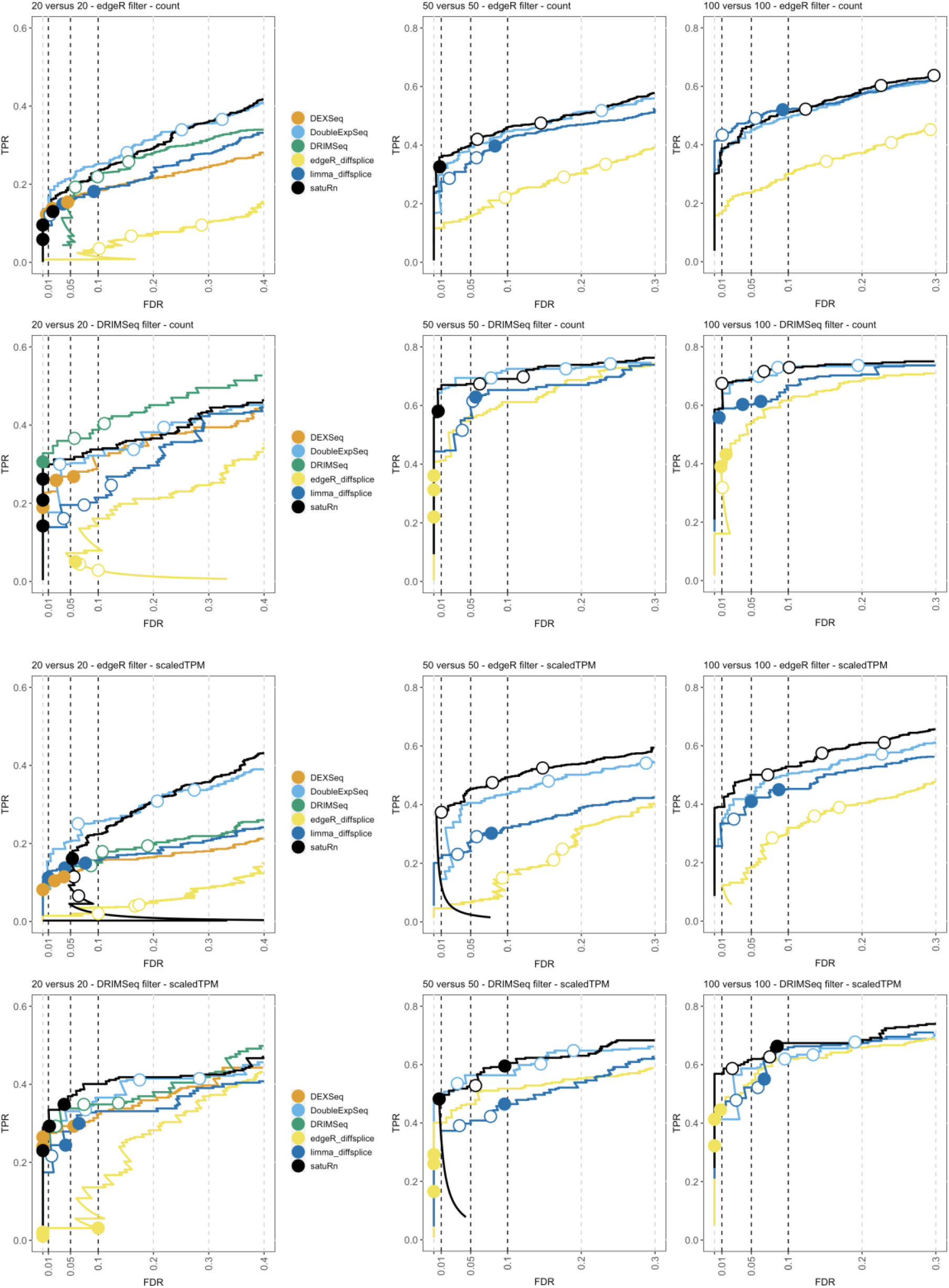
Performance evaluation of satuRn on the real scRNA-Seq dataset by Darmanis et al. FDR-TPR curves visualize the performance of each method by displaying the sensitivity of the method (TPR) with respect to the false discovery rate (FDR). The three circles on each curve represent working points when the FDR level is set at nominal levels of 1%, 5% and 10%, respectively. The circles are filled if the empirical FDR is equal or below the imposed FDR threshold. We generated three two-group comparisons of 20, 50 and 100 cells each (left, middle and right panel, respectively). The benchmark was performed both on the raw counts **(rows 1 and 2)** or on scaled transcripts-per-million (TPM) **(rows 3 and 4)** as imported with the Bioconductor R package tximport^59^. We additionally adopted two different filtering strategies; an edgeR-based filtering **(rows 1 and 3)** and a DRIMSeq-based filtering **(rows 2 and 4)**. Overall, the performance of satuRn is similar to DoubleExpSeq, the best tools from the literature. In addition, our method consistently controls the FDR close to its imposed nominal FDR threshold, while DoubleExpSeq becomes more liberal with increasing sample sizes. On the dataset with the smallest sample size, the FDR control of *satuRn* does become too strict.

**Figure S10:**
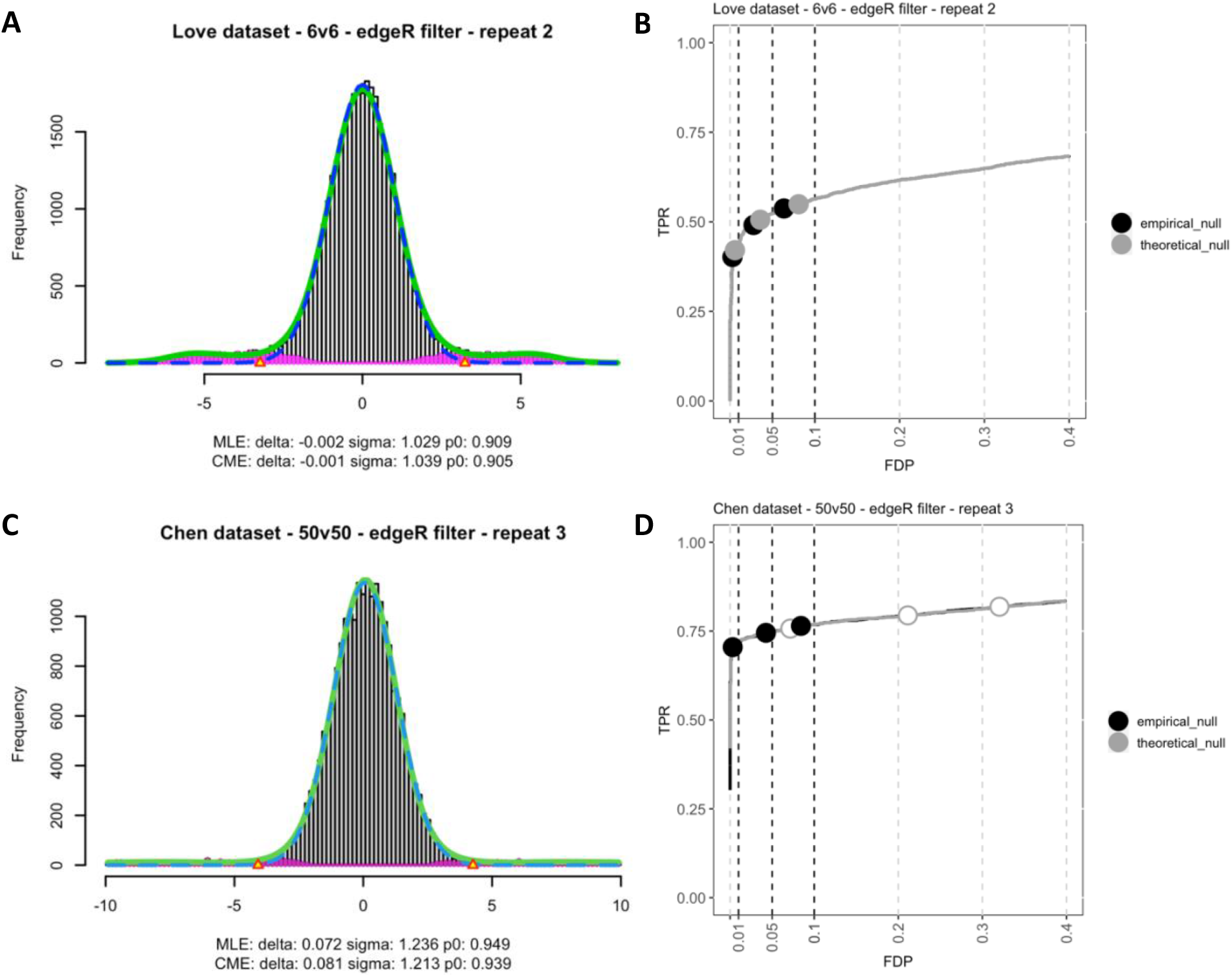
The effect of using an empirical null distribution on the false discovery control of satuRn. **Panel A:** Empirical distribution of the satuRn test statistics in one of the bulk transcriptomics benchmark datasets adapted from Love *et al*. The test statistics are z-scores, calculated from satuRn p-values as described in formula 5 (see Methods). As this benchmark dataset is constructed to have 15% DTU transcripts and thus 85% non-DTU or null transcripts, most of these z-scores are expected to follow a standard normal distribution (mean = 0, standard deviation = 1). This is reflected in the maximum likelihood estimates for the mean and variance of the empirical null distribution (mean = − 0.002, standard deviation = 1.029). **Panel B:** Corresponding FDP-TPR curve for the bulk transcriptomics benchmark dataset. As the theoretical null distribution and the empirical null distribution are virtually identical, we observe a negligible difference between both strategies, both in terms of performance and FDR control. **Panel C:** Empirical distribution of the satuRn test statistics in one of the single-cell benchmark datasets adapted from Chen *et al*. Again, most of these z-scores are expected to follow a standard normal distribution as this benchmark dataset is also constructed to have 15% DTU transcripts and thus 85% non-DTU or null transcripts. However, the empirical distribution is considerably wider than expected (standard deviation = 1.236). We additionally observe a small shift of the distribution (mean = 0.072). **Panel D:** Corresponding FDP-TPR curve for the single-cell benchmark dataset. While the inference for satuRn is overly liberal when working under the theoretical null, FDR control is restored by adopting the wider empirical null distribution. Note that the performance will only be affected when the empirical null distribution is strongly shifted with respect to the theoretical null (i.e., a large mean in absolute value), which was not the case in this example nor in any other dataset from our analyses.

**Figure S11:**
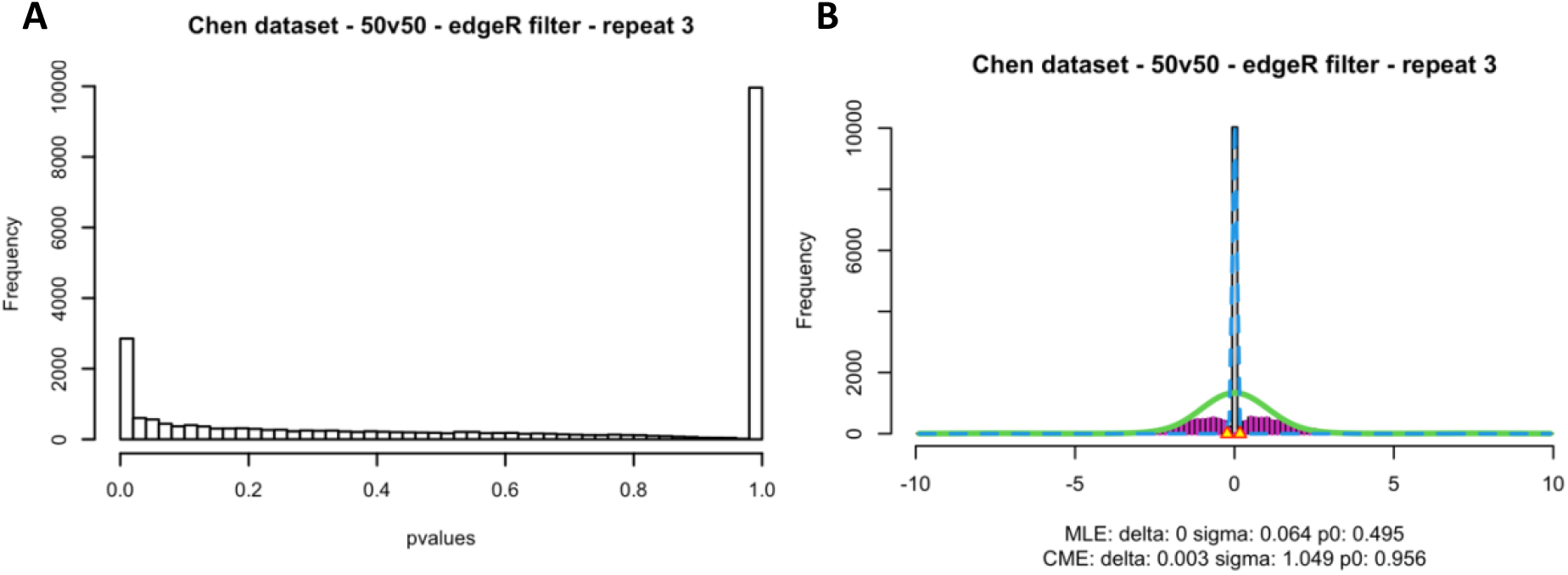
Adopting an empirical null distribution to improve FDR control is infeasible for DoubleExpSeq. **Panel A:** Distribution of the p-values from a DoubleExpSeq analysis in one of the single-cell benchmark datasets adapted from Chen *et al*. We immediately observe the large spike of p-values equal to 1, which distorts the p-value distribution. In addition, the p-values in the mid-range (e.g., from 0.1 to 0.9), which are expected to be uniformly distributed, are skewed towards smaller values, which underlies the overly liberal results of DoubleExpSeq in our single-cell benchmarks. **Panel B:** The corresponding empirical distribution of the DoubleExpSeq test statistics. The test statistics are z-scores, calculated from the original DoubleExpSeq p-values as described in formula 5 (see Methods). As all our benchmark datasets are constructed to have 15% DTU transcripts and thus 85% non-DTU or null transcripts, most of these z-scores are expected to follow a standard normal distribution (mean = 0, standard deviation =1). However, given the pathological distribution of the p-values it is not feasible to properly estimate the empirical null distribution, as also clearly shown by the widely different parameter estimates obtained using the two estimation frameworks implemented in the *locfdr* R package; compare the estimates between MLE (maximum likelihood estimation) and CME (central matching estimation).

**Figure S12:**
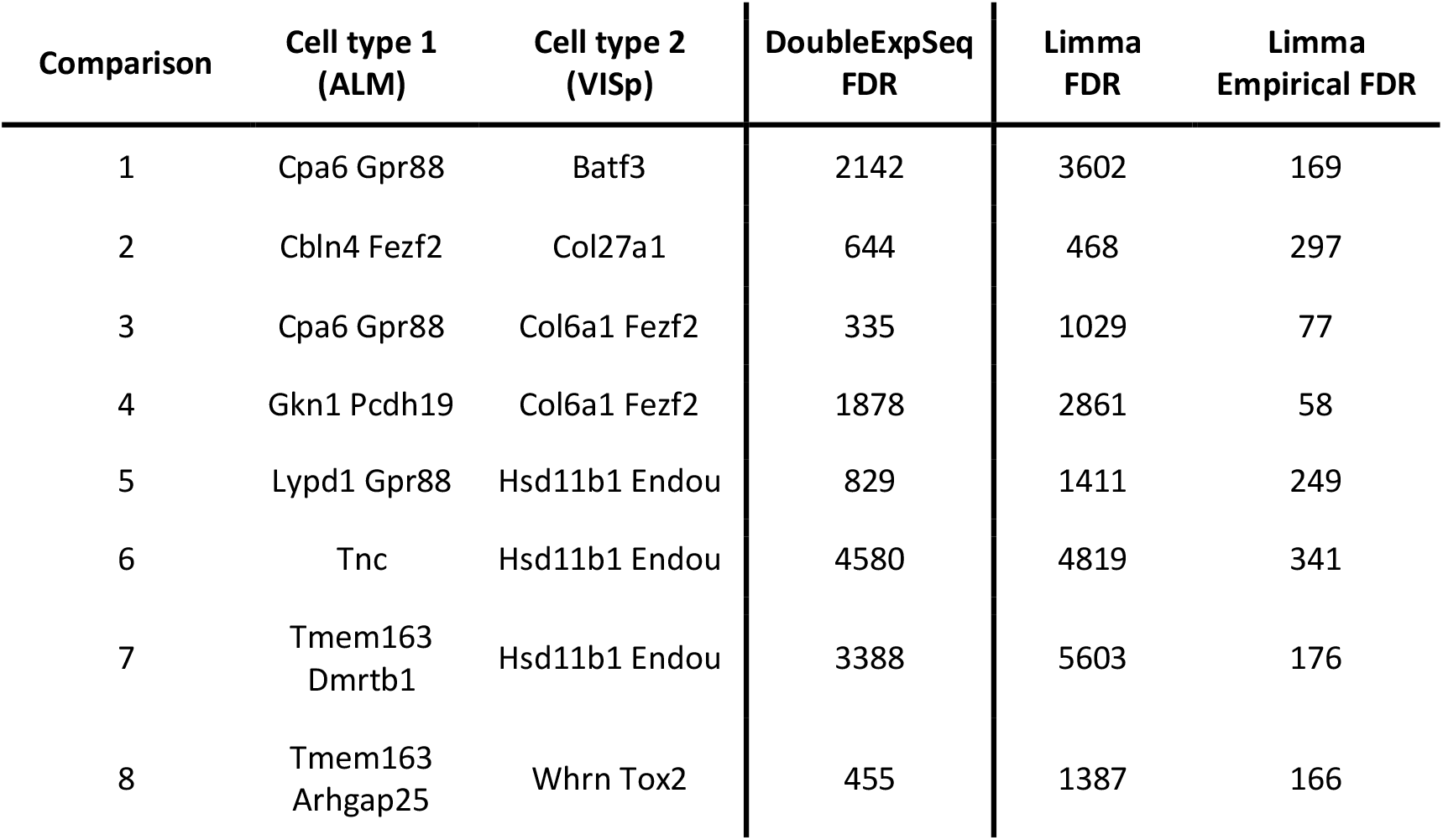
Number of differentially used transcripts as identified by DoubleExpSeq and limma diffsplice. The first three columns indicate the comparisons between ALM cell types (column 2) and VISp cell types (column 3), respectively. Column 4 indicates the number of differentially used transcripts as identified by DoubleExpSeq. Column 5 indicates the number of differentially used transcripts as identified by a limma diffsplice analysis with default settings. Column 6 displays the number of differentially used transcripts found by limma diffsplice after correcting for deviations between the theoretical and empirical null distributions.

**Figure S13:**
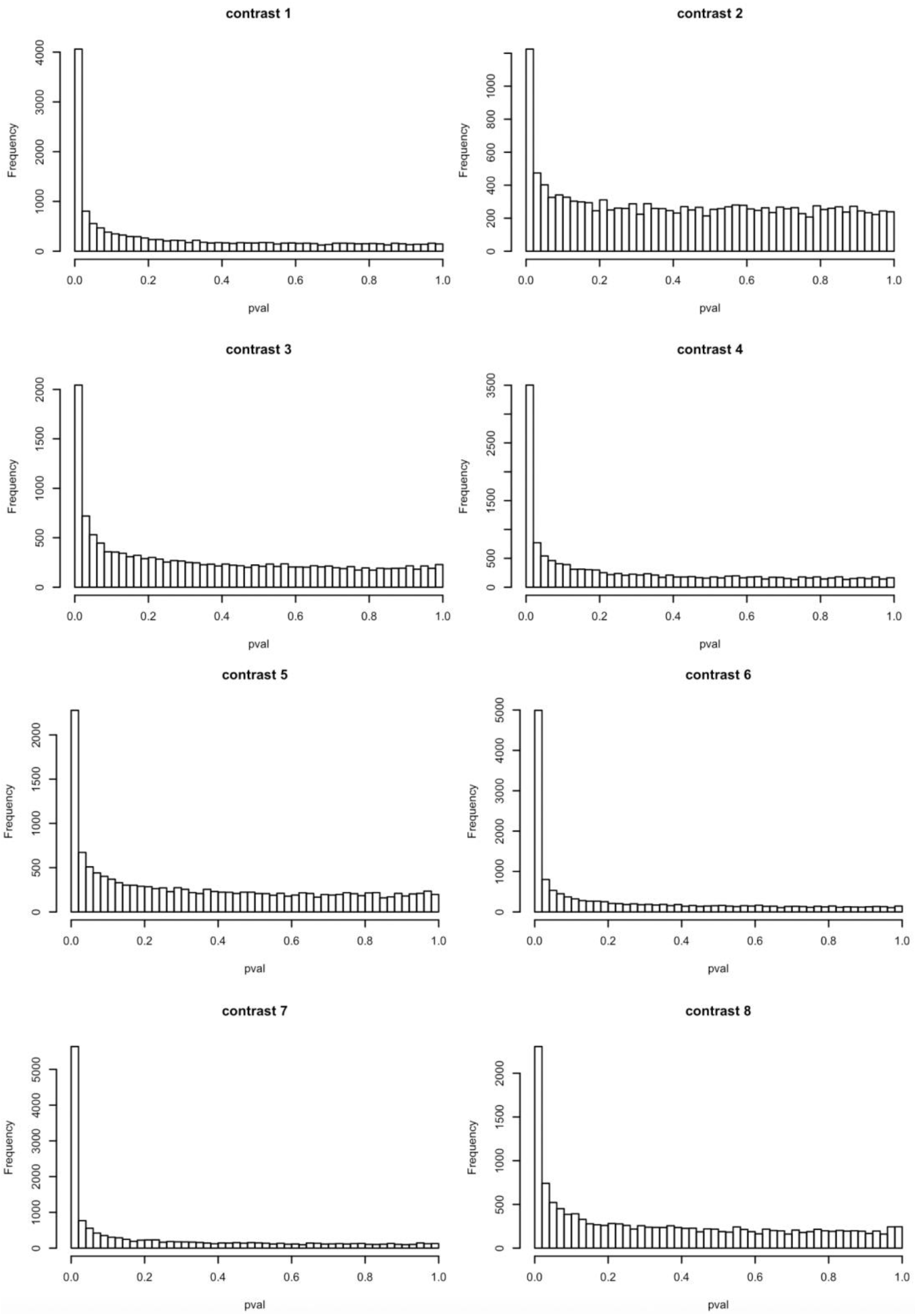
Histograms of the p-values from limma diffsplice. From these histograms, the huge number of DTU transcripts identified by limma diffsplice become apparent. Note that the general tendency of limma diffsplice for smaller p-values is better visible when converting the p-values into z-scores (see Figure S13)

**Figure S14:**
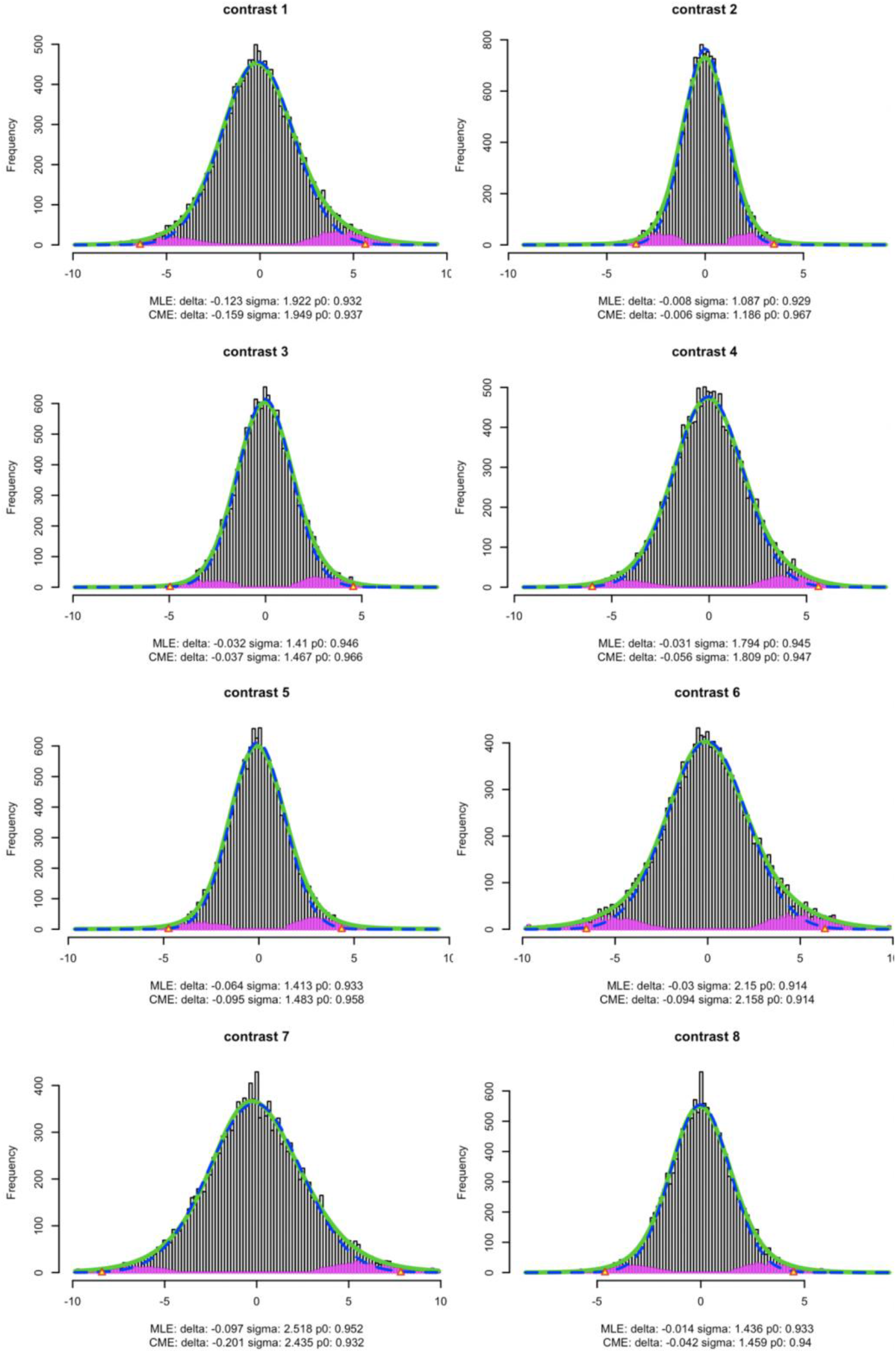
Empirical distribution of the limma diffsplice test statistics. The test statistics are z-scores, calculated from limma diffsplice p-values as described in formula 5. Theoretically, these z-scores are expected to follow a standard normal distribution (mean = 0, standard deviation =1). Here, however, the empirical distributions are considerably wider (standard deviation > 1), as indicated underneath the plots. This indicates that the results returned by limma diffsplice in this case study are overly liberal.

**Figure S15:**
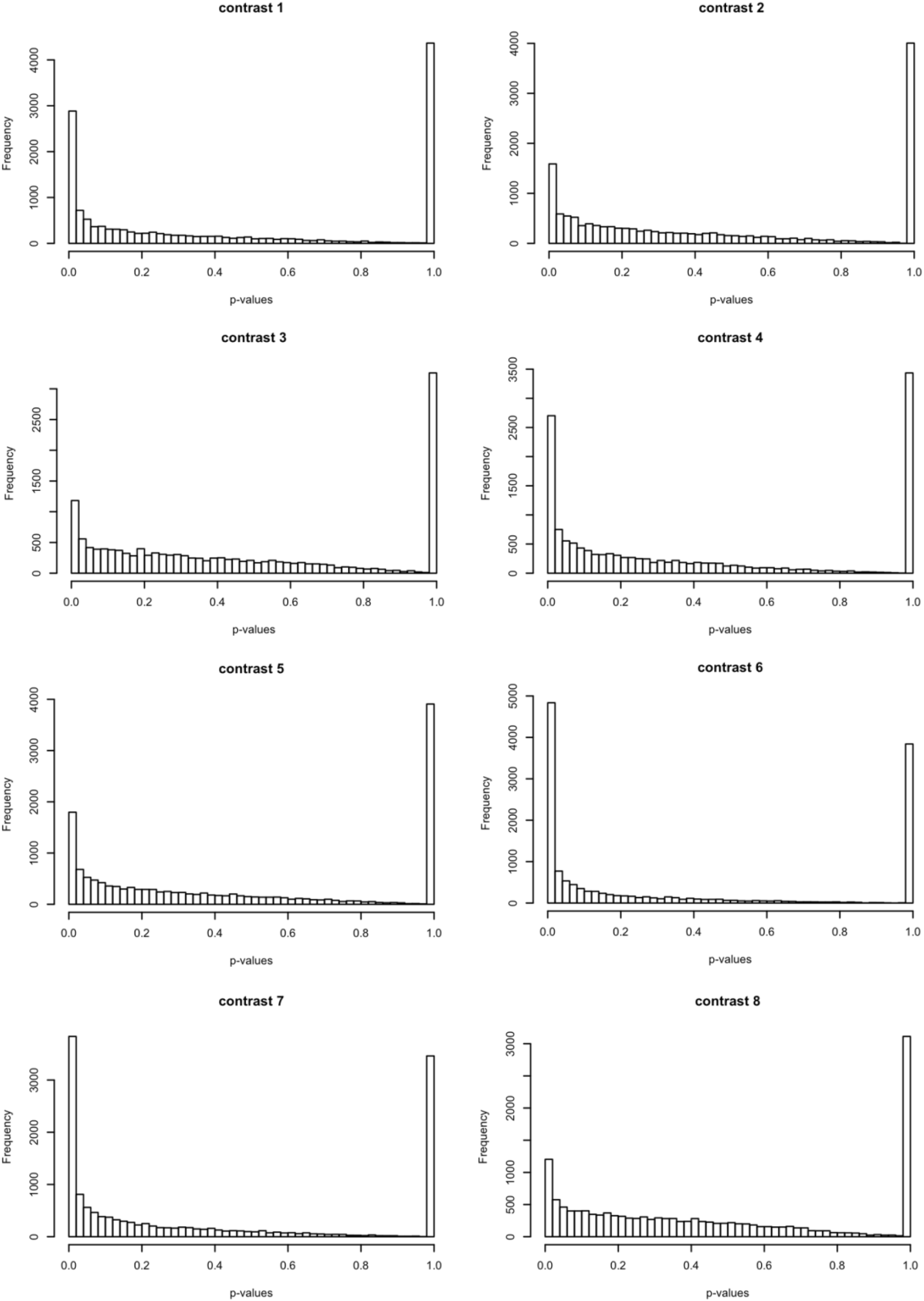
Histograms of the p-values from DoubleExpSeq. From these histograms, the huge number of DTU transcripts identified by limma diffsplice become apparent. In addition, we observe a gradual decrease of p-values over the interval [0.05 < p < 0.95], with a very large spike of p-values that are exactly 1 in all comparisons or contrasts of interest.

**Figure S16:**
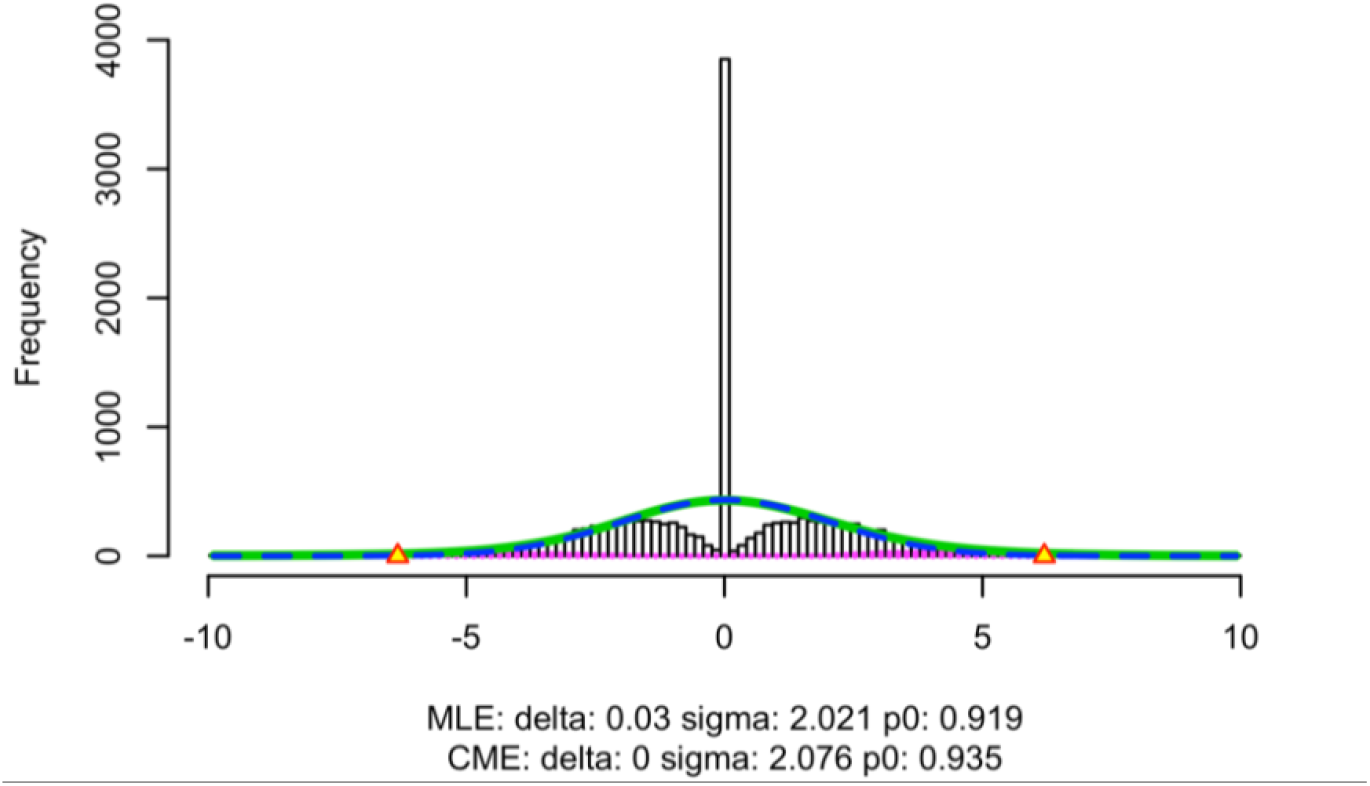
Empirical distribution of the test statistics in comparison #6 of the case study with DoubleExpSeq. The test statistics are z-scores, calculated from DoubleExpSeq p-values as described in formula 5 (see Methods). Theoretically, the bulk of these z-scores are expected to follow a standard normal distribution (mean = 0, standard deviation =1), i.e., assuming that most transcripts are not differentially used. However, the large spike of p-values equal to 1 (See Figure S14) results spike of z-scores equal to zero, which poses a problem when estimating the empirical null distribution (blue dashed curve).

